# Enrichment-free glycoproteomics harnessing real-time mass defect-driven glycopeptide classification reveals sex differences in murine fucosylation

**DOI:** 10.64898/2026.07.03.736455

**Authors:** Bingyuan Zhang, The Huong Chau, Kristina Mae Bienes, Hiromu Arakawa, Masaya Hane, Chihiro Sato, Akira Yokoi, Hiroyuki Kaji, Christopher Ashwood, Yusuke Matsui, Rebeca Kawahara, Morten Thaysen-Andersen

## Abstract

Glycopeptide enrichment remains a cornerstone in glycoproteomics, but bias and reproducibility issues continue to hinder biological insight and clinical translation. Employing curated glycoproteomics datasets and machine learning, we trained a glycopeptide classifier to recognize N-glycopeptide precursors through mass defect signatures. Integration of the classifier into a data-dependent acquisition framework facilitated real-time prediction of N-glycopeptides from human serum and revealed sex differences in murine plasma fucosylation opening avenues for enrichment-free glycoproteomics.

---

Protein glycosylation impacts most biological processes and pathological conditions positioning glycoproteins as a largely untapped source of disease biomarkers and therapeutic targets^1,2^. However, glycopeptide profiling from biological specimens remains challenging^3^. Consequently, the vast glycoproteome is still poorly mapped across pathophysiological conditions and methodological bottlenecks continue to prevent clinical translation^4^.

Given the suppression of glycopeptide ions by their non-glycosylated counterparts^5^, current glycoproteomics workflows still rely on glycopeptide enrichment before LC-MS/MS detection. Asn-linked (N-) glycoproteins, most notably, display extensive site heterogeneity causing further signal suppression^6^. Despite key advances^7^, glycopeptide enrichment methods remain inefficient, labor-intensive and often biased to specific glycoforms. Consequently, glycopeptides are commonly underrepresented, quantitively skewed or completely missed using traditional intensity-based data-dependent acquisition (DDA) adversely impacting the glycoproteome coverage and hindering biological and clinical insight^8^.

As noted previously^9^, N-glycopeptides have proportionally more oxygen (15.9949 Da) and less nitrogen (14.0031 Da) than non-glycosylated peptides, **Figure 1a**. In a global analysis (**Online Methods**), glycopeptides from both a simulated human N-glycoproteome (**Supplementary Figure S1)** and our own experimental datasets indeed featured profoundly raised oxygen and reduced nitrogen levels relative to unmodified peptides, **Figure 1b** and **Supplementary Figure S2**. Consistently, N-glycopeptides exhibited mass defect (MD) signatures that were distinctly different from non-glycosylated peptides, **Figure 1c**. Thus, MD represents a universal yet under-utilized molecular trait to distinguish N-glycopeptides in mixtures.

**Figure 1.**
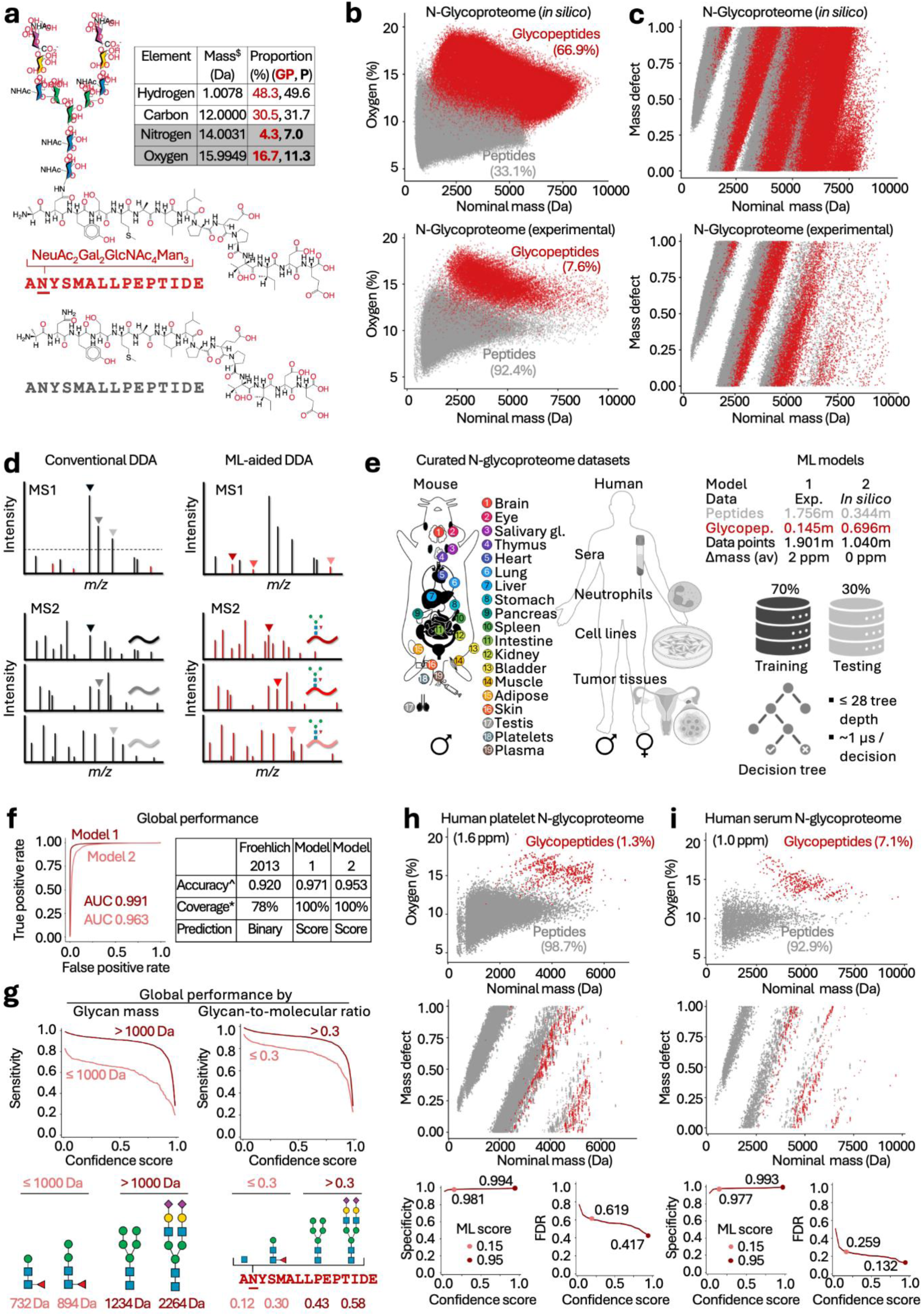
MD-driven glycopeptide classifier. **a**) Oxygen (negative MD) and nitrogen (positive MD) content of a representative N-glyco/peptide pair. ^$^Monoisotopic mass. **b-c**) Global oxygen levels and MD-to-nominal mass plots of all tryptic glyco/peptides from simulated (top) and experimental (bottom) N-glycoproteome data. See **Supplementary Figure S1** for details of the simulated N-glycoproteome and **Supplementary Figure S2** for matching nitrogen plots. **d**) Concept of the ML-aided glycopeptide classifier. **e**) Decision tree-type models were trained on curated glycoproteomics data (model 1, see **Supplementary Table S1**) or from simulated N-glycoproteome data (model 2). **f**) Data points (30%) were held out for benchmarking against a simple binary glycopeptide classifier covering only a limited *m/z* range (*)^9^. ^Accuracy calculated based on ML0.95 score cut-off. Global performance of model 1 by N-glycan mass (left) and glycan-to-molecular mass ratio (right) with examples provided for context (bottom). Dataset-specific oxygen levels, MD-to-nominal mass plots and prediction accuracy for N-glycoproteome data of **h**) human platelets and **i**) human serum previously unseen by the ML model (see **Supplementary Table S2)**. See **Supplementary Figure S3** for performance assessed on datasets used for training.

Based on this principle, we sought to establish a real time-compatible glycopeptide classifier that can augment conventional intensity-based DDA approaches by selecting, with precision, N-glycopeptide precursor ions for fragmentation based on MD signatures rather than by intensity, **Figure 1d**. Opting for an explainable, confidence-based and prompt (1 µs/prediction) decision tree strategy, we trained and tested two ML models using either curated N-glycoproteomics datasets of diverse human and mouse tissues (∼1.9m data points, model 1) or simulated N-glycoproteome data (∼1.0m data points, model 2), **Figure 1e** and **Supplementary Table S1**. Model 1 (AUC 0.991) showed comparably higher global performance and consistently outperformed a simple (binary) MD-based glycopeptide classifier^9^, **Figure 1f**. The experimentally-trained model 1 was therefore used as our preferred glycopeptide classifier.

Intuitively, the ML model performed better for peptides carrying relatively large N-glycans (≥6-7 monosaccharide residues) featuring relatively high glycan-to-molecular mass ratios (>0.3), **Figure 1g**. The glycopeptide classifier showed higher performance (lower FDRs) for serum known to harbor glycoproteins with relatively large N-glycans^10^ (averaging ∼2,100 Da) than for other tissue specimens used in the training data (typically 1,200-1,800 Da), **Figure 1h-i, Supplementary Figure S3** and **Supplementary Table S2**. While these attributes make the glycopeptide classifier ideal for serum and plasma N-glycoproteomics, modest performance is expected for peptides with low glycan-to-molecular mass ratios (e.g. short O-glycopeptides).

To show proof-of-concept for enrichment-free serum N-glycoproteomics using the ML-aided DDA method, we tested glycopeptide prediction in both off-line (static) and on-line (real-time) setups and compared these against conventional DDA, **Figure 2a-b**. Taking less than 0.1 milliseconds for up to 100 predictions per MS1 survey scan, the real-time glycopeptide prediction added an insignificant time penalty to the 3 s duty cycle. Both off- and on-line ML-aided DDA outperformed conventional intensity-based DDA in terms of glycoPSM counts and proportion of glycopeptide spectra acquired, **Figure 2c-d**. Increasing the confidence threshold of the classifier from ML0.15 to ML0.95 markedly improved the prediction accuracy as shown by higher glycopeptide-to-peptide ratios without compromising the glycoproteome coverage.

**Figure 2.**
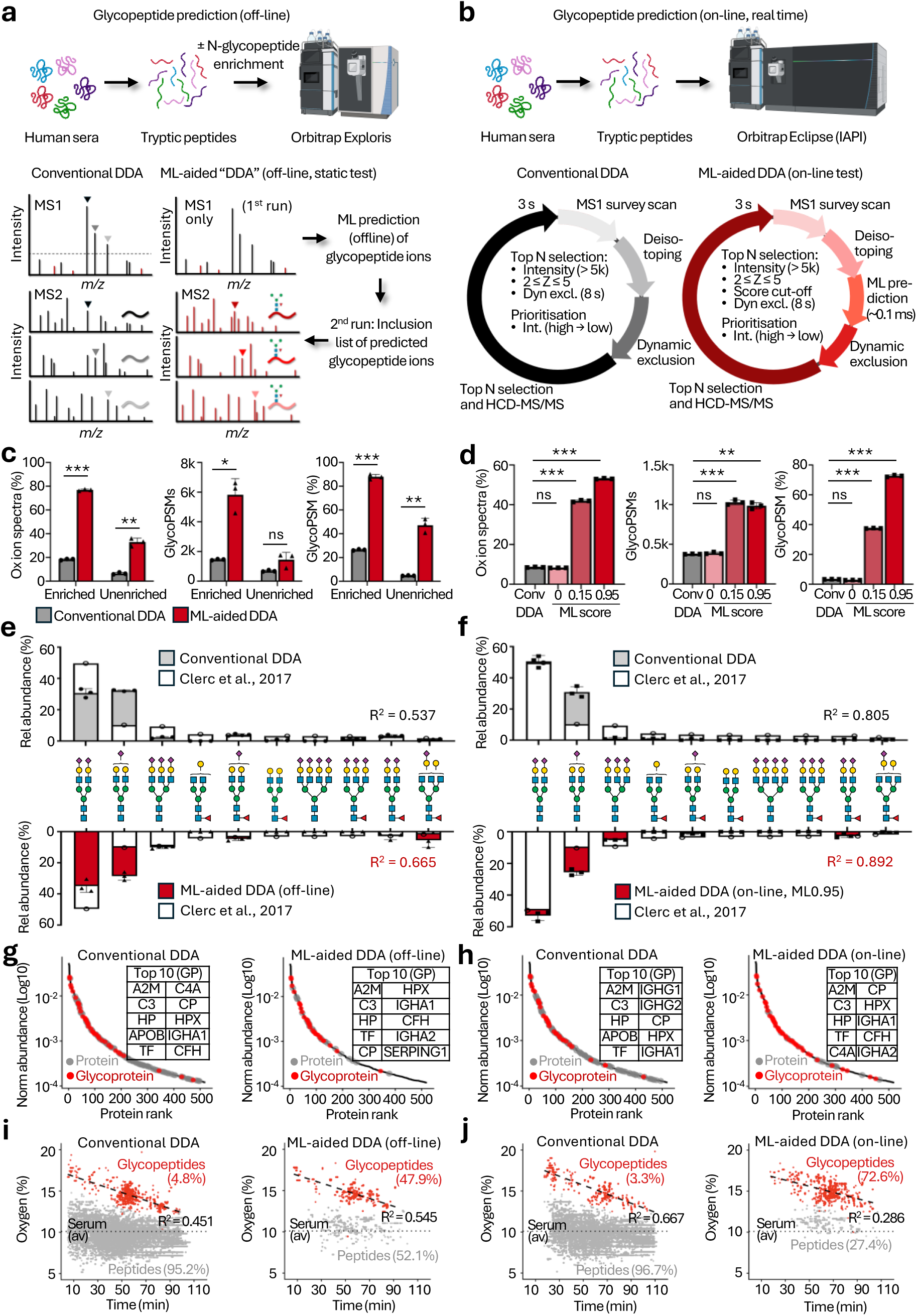
Enrichment-free serum glycoproteomics. The glycopeptide classifier was tested head-to-head against conventional intensity-based DDA on human sera using an **a)** off-line (static) approach employing an inclusion list to validate predicted glycopeptide precursors in a subsequent targeted LC-MS/MS run and **b**) on-line (real-time) approach in which the ML model embedded into the IAPI predicted glycopeptide precursor ions directly from unenriched peptide mixtures. Proportion of oxonium ion spectra (left) and glycoPSMs (right) of all fragment spectra and absolute glycoPSM count (middle) of conventional (grey) and ML-aided (red) DDA performed **c)** off-line and **d**) on-line. Two-tailed, unpaired welch t-tests, \*\*\**p* < 0.001, \*\**p* < 0.01, \**p* < 0.05, ns, not significant. Correlation analyses of the known serum N-glycome^10^ against conventional DDA (top) and ML-aided DDA (bottom) performed **e**) off-line and **f**) on-line. The glyco/proteins identified with conventional DDA and using **g**) off-line and **h**) on-line ML-aided DDA approaches were mapped on the known abundance range of the serum proteome (normalized to 1)^11^. Top glycoproteins are listed. Oxygen levels of glyco/peptides identified with conventional DDA and using **i**) off-line and **j**) on-line ML-aided DDA approaches. Dotted grey line: Oxygen content of typical serum peptides^11^. Black broken line: trendline for glycopeptide data points from ML-aided DDA.

Next, we tested for potential bias of the N-glycopeptides predicted by the ML model. Benchmarking against the known serum N-glycome^10^, both the static and real-time ML-aided DDA experiments showed comparably less N-glycan bias than conventional DDA, **Figure 2e-f**. The identified glycoproteins were, as expected, mostly those known to be of high abundance in serum^11^ including α-2-macroglobulin, complement-3 and haptoglobin, **Figure 2g-h**.

Validating that the classifier is guided by discriminatory MD signatures, the correctly predicted glycopeptides (and the minor fraction of mis-predicted non-glycosylated peptides) exhibited higher oxygen and lower nitrogen levels compared to average serum peptides^11^, **Figure 2i-j** and **Supplementary Figure S4**. Consistent with the observation that short peptides are poorly retained in C18-LC, early eluting glycopeptides exhibited substantially raised oxygen and suppressed nitrogen levels relative to typical tryptic peptides of serum proteins. These observations were recapitulated in the glycan-to-molecular mass ratios, **Supplementary Figure S5**. Collectively, this confirms that MD drives the prediction mechanism of the glycopeptide classifier.

To demonstrate the potential for biological discovery, we applied ML-aided DDA-based glycoproteomics to neat plasma from male and female mice, **Figure 3a**. Consistent with observations from human sera, ML-aided DDA dramatically outperformed conventional intensity-based DDA in terms of identified glycoPSMs and glycoproteins, **Figure 3b**. Notably, the LC-MS/MS data generated by ML-aided DDA indicated separation of male and female plasma, trends that were not apparent with conventional DDA, **Figure 3c**. Interrogation of the ML-aided DDA data revealed prominent sex differences in fucosylated glycoforms e.g. HexNAc_4_Hex_5_Fuc_1_NeuGc_2_, HexNAc_7_Hex_8_Fuc_1_NeuGc_3_ across various glycoproteins e.g. kinogen-1 (O08677 and corticosteroid-binding globulin (Q06770), **Figure 3d-e**. In line with our recent study reporting that male and female mice differ in FUT8-mediated core fucosylation across specific tissues^12^, the ML-aided DDA data showed considerably raised fucose levels in plasma from female mice, findings that were independently validated by quantitative TMT-DDA of enriched N-glycopeptides from the same samples, **Figure 3f**.

**Figure 3.**
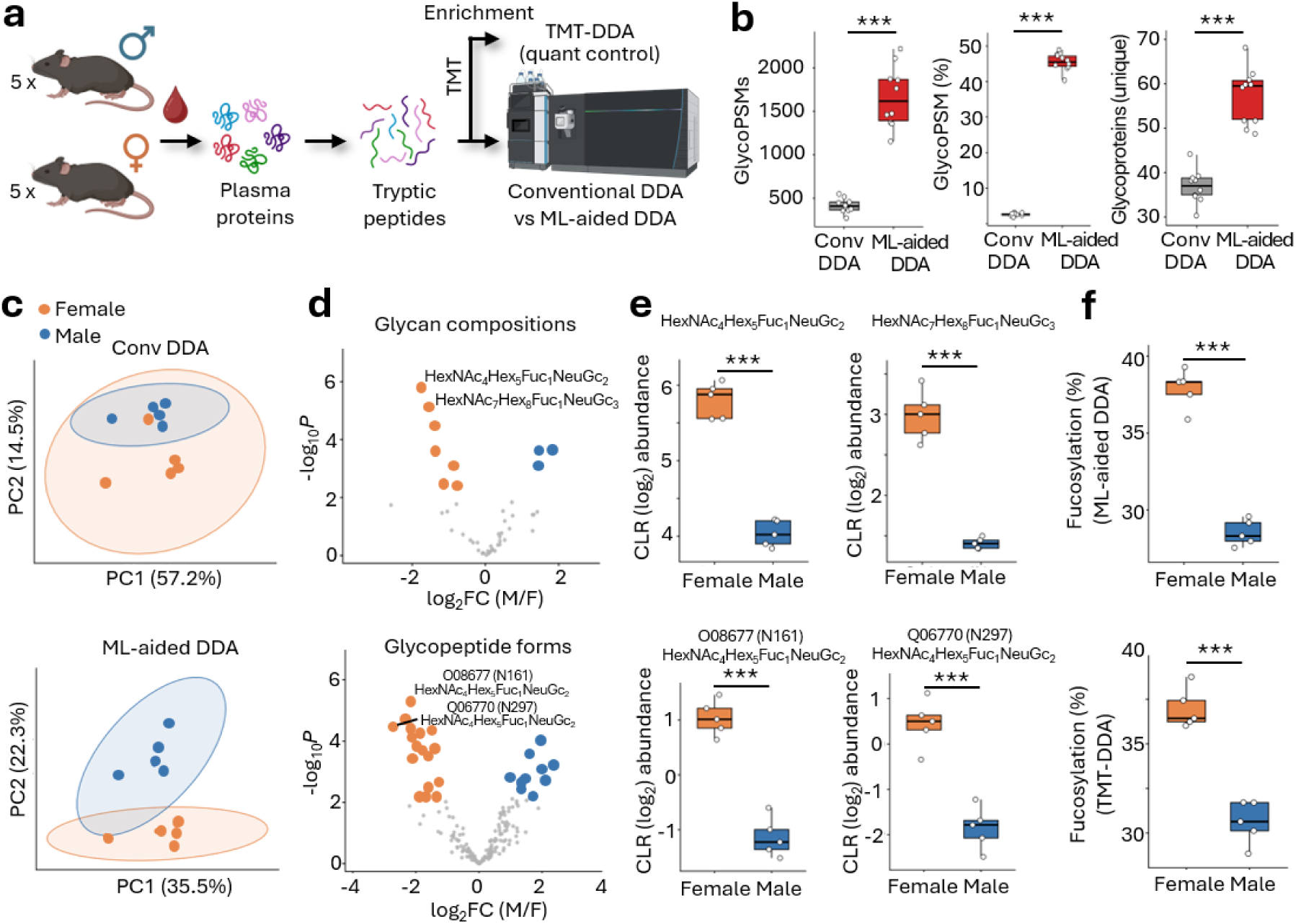
Glycopeptide classifier reveals sex differences in fucosylation in murine plasma. **a**) Unenriched peptide mixtures from plasma proteins of male and female mice (n = 5/sex) were analyzed using conventional DDA and ML-aided DDA as well as after TMT-labelling and glycopeptide enrichment as a quantitative control. **b**) Counts and proportion of glycoPSMs and unique glycoproteins. Paired two-tailed t-tests. **c**) PCA of LC-MS/MS data obtained by conventional DDA (top) and ML-aided DDA (bottom). Interrogating the ML-aided DDA data, **d**) volcano plots and **e**) box plots of glycan compositions (top) and glycopeptide forms (bottom) comparing female (orange) and male (blue) plasma. Moderated two-tailed t-tests. **f**) Global level of glycopeptide fucosylation measured by ML-aided DDA (top) and TMT-DDA (quantitative control, bottom). Welch unpaired two-tailed t-tests. For boxplots (b, d, f), whiskers indicate spread of data and individual data points are shown, \*\*\**p* < 0.001, \*\**p* < 0.01, \**p* < 0.05, ns, not significant.

Glycopeptide enrichment remains central to glycoproteomics workflows, but inherent limitations of this manual and laborious sample handling step continue to hinder biological insight and clinical translation. To address this long-standing issue, we have here shown proof-of-concept for and biological discovery with a real-time compatible classifier that instantly recognizes N-glycopeptide precursor ions from complex MS1 survey scans. By integrating the ML model into a DDA framework, informed selection of glycopeptide precursors for downstream MS/MS sequence analysis were achieved directly from crude biological specimens without the need for prior glycopeptide enrichment. The classifier is conceptually also compatible with TMT-labeled peptides and other proteolytic enzymes such as promiscuous endo-proteases generating short glycopeptides resulting in higher glycan-to-molecular mass ratios even for O-glycopeptides. Conclusively, the glycopeptide classifier, which is vendor-free and, in principle, is compatible with any instrument featuring an open instrument application programming interface (IAPI) advances the field towards enrichment-free glycoproteomics.

## Data availability

The LC-MS/MS datasets used to train the glycopeptide classifier are publicly available through the ProteomeXchange Consortium via the PRIDE repository (PXD074760, PXD064579, PXD065024, PXD056996, PXD039387). Newly generated LC-MS/MS datasets used to demonstrate proof-of-concept and the potential for biological discoveries of the glycopeptide classifier are publicly accessible (PXD080247, token: ghxhUPMyi2Kz and PXD074760).

## Code availability

The code for the glycopeptide classifier is available free of charge.

## Online Methods

### Overview

In short, we trained and benchmarked a real-time compatible glycopeptide classifier, used off-and on-line strategies to show proof-of-concept for enrichment-free glycoproteomics of human sera and demonstrated potential for biological discovery by applying the method to plasma from male and female mice. Using a decision-tree model, the glycopeptide classifier was trained using a collection of curated N-glycoproteomics datasets (model 1) or simulated N-glycoproteome data (model 2). The experimental glycoproteomics datasets were generated from diverse mouse and human tissues (**Supplementary Table S1**) whereas only human N-glycoproteome data was *in silico* simulated (**Supplementary Figure S1**).

### Concept and design

As data input, both models have been developed to use nominal mass (NM) and mass defect (MD) information that can readily be extracted from deconvoluted precursor data from MS1 survey scans of tryptic N-glycopeptides and non-glycosylated peptides. The output from the glycopeptide classifier, in turn, is a confidence-based probability score predicting whether a specific precursor ion (with a given NM and MD) correspond to a tryptic N-glycopeptide or a tryptic non-glycosylated peptide (see Model architecture and training).

Following training and optimization of the glycopeptide classifier, the global performance of the two models was benchmarked using a proportion of data points (30%) held out from the experimental training datasets. Performance of the classifier was evaluated based on several metrics (e.g. AUC, accuracy, FDR, GlycoPSMs, see below). Furthermore, we tested the performance of model 1 on a separate set of N-glycoproteomics datasets not used for training (see Model evaluation and benchmarking) and only this experimentally-trained model was used for the downstream proof-of-concept and application experiments.

For the proof-of-concept experiments, off- and on-line ML-aided DDA was tested against conventional intensity-based DDA using the same sample (tryptic peptide mixture of human sera). Identical LC-MS/MS settings were used across the different setups where possible to enable a fair head-to-head comparison. For the off-line (static) approach, the classifier first predicted glycopeptide precursor ions from MS1 survey scans of LC-MS files acquired without fragmentation, and the predicted N-glycopeptide candidates were then compiled in an inclusion list, which were targeted in a second injection of the same sample, see **Figure 2a** for graphical overview. For the on-line (real-time) testing, the glycopeptide classifier was re-written into C# code and implemented within the instrument application programming interface (IAPI) of the mass spectrometer to enable prediction of glycopeptide precursor ions in real time, see **Figure 2b** for graphical overview.

To demonstrate the potential for biological discovery, we applied the online ML-aided DDA-based glycoproteomics method to protein extracts of plasma from five female and five male mice and compared the outputs with conventional intensity-based DDA, see **Figure 3a** for graphical overview. In doing so, we maintained the same conditions for the glycopeptide classifier and LC-MS/MS settings as those used in the previous proof-of-concept experiments to demonstrate that the ML-aided DDA method can be readily applied across species and specimens to enable new biological discoveries that conventional DDA fail to capture. To quantitatively validate the findings from the ML-aided DDA experiments of murine plasma, the same samples were analyzed after TMT labelling and glycopeptide enrichment.

### Experimental training data for the glycopeptide classifier

Experimental N-glycoproteome data from a total of 234 LC-MS/MS files acquired from 19 mouse tissues and four human sample types of different (patho)physiological conditions (e.g. normal, sepsis, cancer…) were used for the training and benchmarking of the glycopeptide classifier (see **Supplementary Table S1** for overview of tissues and data files). Our curated collection of N-glycoproteomics datasets has been published and are publicly available^12–17^.

To obtain a balance of glycopeptides and non-glycosylated peptides, the selected LC-MS/MS files contained data acquired from both enriched glycopeptide samples and peptide mixtures analyzed without enrichment. Datasets were chosen to maximize the diversity of glycoproteome training data to generate an all-round and well-balanced N-glycopeptide classifier rather than focusing on a specific tissue or species. Only datasets of unlabeled tryptic glyco/peptides were considered for the training of the classifier, and all training files were acquired using intensity-based DDA approaches with Thermo Orbitrap LC-MS/MS instrumentation to mimic the downstream proof-of-concept testing and biological application performed on the same type of equipment. As MS1 accuracy (and in turn MD) is critical to the glycopeptide classifier, only datasets with high mass accuracy (< 10 ppm) were considered noting that the actual delta mass was ∼2 ppm (average) for all glycopeptides and non-glycosylated peptides used for the training of the glycopeptide classifier.

To generate reliably labelled data points for the model training, glycopeptides (glycoPSMs) and non-glycosylated peptides (non-glycoPSMs, alternatively written as PSMs) were confidently identified directly from the selected LC-MS/MS raw files. For this purpose, Byonic v5.1.1 (Protein Metrics) was used to search the mouse tissue glycoproteome data, and human ovarian tissue and extracellular vesicle samples, while Byonic v5.6.16 was used to search the human neutrophil samples and Byonic v.4.5.2 used to search human and mouse platelets and human sera. Searches were performed against either the human proteome database (*homo sapiens*; 20,360 entries, UniProtKB, November 2021) or the murine proteome database (*mus musculus*; 17,202 entries, UniProtKB, May 2024), depending on the sample origin.

Sample-specific N-glycan search spaces were used for glycopeptide identification. For human neutrophil samples, a glycomics-informed search space comprising 47 N-glycan compositions was used. For mouse tissue LC-MS/MS files used for training, searches were performed using a default mammalian N-glycan database containing N-glycan compositions without sodium adducts, with the paucimannosidic composition M3F (Man_3_GlcNAc_2_Fuc_1_) manually added, see **Supplementary Table S3**. To search the LC-MS/MS files of human samples used for training, NeuGc-containing compositions were manually removed from the glycan search space to minimize incorrect assignments arising from isobaric or near-isobaric glycan compositions, see **Supplementary Table S4**.

All searches were performed using precursor and fragment ion mass tolerances of 10 ppm and 20 ppm, respectively. Carbamidomethylation of cysteine residues (+57.021 Da) was set as a fixed modification. Fully tryptic peptides were searched, allowing up to two missed cleavages per peptide. Methionine oxidation (+15.994 Da) was defined as a common variable modification, while N-glycosylation of sequon-localized asparagine residues was specified as a rare modification. Variable modifications were limited to a maximum of two common modifications and one rare modification per peptide.

Confidence was achieved by consistently applying stringent scoring thresholds (PEP2D < 0.001) across all searches of the training data. While our datasets comprised nearly 5.0m spectral data points, these were filtered to ∼1.9 million data points by removing isobaric and near-isobaric precursors with the same labels to avoid redundancy and bias in the training data. The actual data points used for the experimental training of the glycopeptide classifier (model 1) comprised 145k N-glycopeptides (glycoPSMs) and 1.76m non-glycosylated peptides (non-glyco PSMs).

### Simulated N-glycoproteome data for the glycopeptide classifier

To complement the experimental glycoproteomics data, we simulated the human N-glycoproteome. Human proteins reported to carry N-glycosylation (5,810 glycoproteins for *homo sapiens* in UniProtKB) were *in silico* reduced and alkylated with iodoacetamide (+57.07 Da) and then digested with trypsin (R/K↓x, x ≠ P) allowing up to one missed cleavage, **Supplementary Figure S1**. This initial step produced a total of 588,016 human tryptic peptides. Focusing only on MS-friendly peptide sequences between 7 and 60 residues (390,951 peptides), tryptic peptides that contained at least one potential N-glycosylation site (NxS/T, x ≠ P) were considered to be potentially modified with N-glycosylation (46,385 putative N-glycosylated peptides); the remaining population were treated as non-glycosylated peptides (344,566 peptides).

Guided by N-glycan compositional data extracted from the experimental N-glycoproteomics datasets used to train the glycopeptide classifier (see Experimental training data for the glycopeptide classifier), realistic N-glycans were *in silico* installed on each N-glycosylation site of the simulated human N-glycoproteome. For this purpose, the frequency of each N-glycan detected in the experimental datasets was determined and used to sample glycans. For every N-glycosylation site, a total of 15 N-glycans were selected from the “observed N-glycome” using a frequency weighting to mimic the natural site micro-heterogeneity observed for mammalian N-glycoproteins. This approach yielded 696,165 (putative) tryptic N-glycopeptides. Considering also the 344,566 non-glycosylated tryptic peptides, the simulated human N-glycoproteome comprised around 1.04 million data points.

### Model architecture and training

The glycopeptide classifier was trained using a decision tree (a CART model, trained with the rpart method, caret R package). From NM and MD input values, easily computed from MS1 survey scan information, the classifier has been designed to return a confidence score indicating the likelihood that an unknown precursor ion corresponds to a tryptic N-glycopeptide. For the glycopeptide classifier, we chose a decision tree model due to superior speed and interpretability thus making this model compatible with LC-MS/MS implementation (within the IAPI) and real time prediction. Notably, both ordinary decision tree and boosted tree models were established and evaluated in the early phases of the project. However, as the boosted tree approach did not enhance the overall performance relative to an ordinary decision tree model and is more challenging to interpret and thus implement, the boosted tree model was not pursued further.

Two decision tree models were established using different training data; Model 1 was trained on experimental N-glycoproteomics data (∼1.90 million points) and model 2 on simulated N-glycoproteome data (∼1.04 million points, see above for details). For both models, 70% of the data points were randomly selected from the datasets and used for training while the remaining 30% were held out for global performance testing and benchmarking.

The two decision tree models were tuned with five-fold cross-validation. The tree complexity parameter that resulted in the best performance was kept. The tree’s complexity parameter was tested over the values 1, 3, 5, 10, 20 and 50 divided by the number of training points. Each tree returns a probability at its leaves, which were used as an adjustable confidence score in the downstream glycopeptide prediction. As such, a given precursor was classified as a tryptic N-glycopeptide if the confidence score exceeded a defined threshold such as, for example, 0.15 (named ML0.15 in the study) or 0.95 (ML0.95). When the ML score cut-off was set to 0 (ML0), the approach mimicked the conventional intensity-based DDA workflow, a condition used as an instrument control. Accordingly, confidence scores equal to or below the threshold were classified as non-glycosylated tryptic peptides.

### Global and dataset-specific evaluation and benchmarking

Global performance was tested for both model 1 and 2 using 30% of the data held out from the training. For each model, we measured the area-under-the-curve (AUC), sensitivity, specificity, false discovery rate (FDR) and the accuracy at a 0.95 confidence score cut-off (ML0.95) and separately benchmarked the models against a simple binary glycopeptide classifier, which predict candidates over a limited mass range (1300 – 5001 Da)^9^.

Using Byonic-based search results as a guide (see above for search settings), the number of true positive (TP), true negative (TN), false positive (FP) and false negative (FN) predictions could be determined. This, in turn, allowed the following performance metrics to be calculated:

Sensitivity (i.e. the true-positive rate) measures the proportion of genuine positives that were correctly identified:

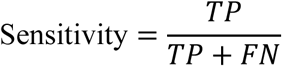

Specificity (i.e. the true-negative rate) measures the proportion of genuine negatives that were correctly rejected:

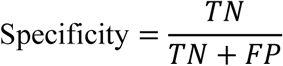

Accuracy measures the overall proportion of correct assignments across both classes:

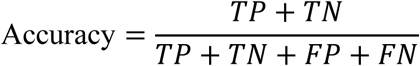

The false discovery rate (FDR) measures the proportion of N-glycopeptides that were incorrectly called:

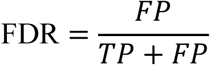

Dataset-specific performance of model 1 and model 2 was first evaluated using the datasets employed for training by holding out 30% of data points. Given its higher global performance, the experimentally-trained model 1 was also separately tested on N-glycoproteomics datasets not used for the training. In both cases, the dataset-specific performance was evaluated using the same performance metrics as for the global performance tests (sensitivity, specificity, accuracy, FDRs).

### Proof-of-concept for enrichment-free serum glycoproteomics

Model 1 was used to document proof-of-concept for enrichment-free serum glycoproteomics.

In preparation for this, protein extract from human sera (200 μg) was reduced with 10 mM dithiothreitol (DTT) for 30 min at 30°C and alkylated with 20 mM iodoacetamide (IAA) for 30 min in the dark at room temperature. Pooled sera were obtained from ovarian cancer patients with permission from Nagoya University HREC (2017–0053). Excess IAA was quenched by the addition of DTT to a final concentration of 20 mM. Samples were digested with sequencing-grade modified porcine trypsin (Promega) at an enzyme-to-substrate ratio of 1:50 (w/w) for 17 h at 37°C. Digestion was terminated by acidification with trifluoroacetic acid (TFA) to a final concentration of 1% (v/v). The resulting peptides were desalted using Oligo R3-C18 SPE micro-columns as described previously (1).

For analysis of unenriched human serum peptides, a 200 μg peptide digest was dried by vacuum centrifugation and reconstituted in 200 μL 0.1% (v/v) formic acid. Subsequently, 1 μL was injected for LC–MS/MS analysis using either conventional intensity-based or ML-aided DDA. For the analysis of enriched N-glycopeptides, the 200 μg serum peptide digest was subjected to hydrophilic interaction liquid chromatography SPE using micro-columns packed with zwitterionic ZIC-HILIC resin (10 μm particle size, 200 Å pore size; kindly provided by Merck Millipore) and C8 disks (Empore) in p10 pipette tips as described previously^2^. The enriched N-glycopeptides were dried by vacuum centrifugation, reconstituted in 20 μL 0.1% (v/v) formic acid, and 4 μL was injected for LC–MS/MS analysis using either intensity-based or ML-aided DDA. Both the enriched and unenriched fractions were used for off-line (static) proof-of-concept experiments, whereas only the unenriched fraction was used for the on-line proof-of-concept experiments.

For the off-line proof-of-concept experiments, the glycopeptide classifier (model 1) was first applied to data from LC-MS runs acquired exclusively using MS1 survey scans to build a target list of candidates for downstream evaluation of the prediction accuracy. For this purpose, raw LC-MS files were converted to mzML files using ThermoRawFileParser V1.5.0^18^. Because of the restricted size of an inclusion list, we reduced the number of predicted glycopeptide candidates by filtering the mzML files for precursor charge state and intensity. Specifically, only candidates with (i) charge states of 2–5 and (ii) intensities ≥ 1 × 10⁶, or ≥ 5 × 10⁵ and exceeding 1% of the base peak intensity in the MS1 scans were retained. Candidates with confidence scores above 0.99 (ML0.99) were retained. Depending on their observed LC retention time from the MS1 runs, the selected candidates were then divided into 110 one-minute retention time bins. For each bin, all MS1 precursors were kept if the candidates did not exceed 300, otherwise only 300 precursor ions were randomly sampled, with the sampling probability weighted according to peak intensity. In total, ∼20,000 glycopeptide candidates distributed across the 110 bins were targeted by a segmented inclusion list (1 min bins). In our experience, exceeding this volume of candidates resulted in software interface failure. The off-line proof-of-concept experiments were repeated three times for each setup generating technical triplicates. For each technical replicate, we first acquired the MS1 data (i.e. performed a LC-MS run), predicted glycopeptide candidates for the inclusion list, performed a targeted DDA analysis (i.e. a LC-MS/MS run of a second sample injection) using the generated inclusion list, and then for comparison conducted a conventional intensity-based DDA run of the same sample.

For the on-line proof-of-concept experiments, model 1 was employed to predict glycopeptide precursor ions during acquisition, which were then prioritized for downstream MS/MS sequence analysis. To implement the trained model onto the LC-MS/MS instrument, the trained model was converted from R into native C# code: a simple script walked the tree and wrote out an equivalent set of nested if/else rules. We verified that the C# implementation generated the same prediction scores as the original R code. The average prediction speed was approximately 1 μs per precursor ion (∼0.1 ms for 100 predictions), demonstrating compatibility with real-time instrument control. The model was embedded within the Thermo Fisher Instrument API (IAPI) through Helios^19^, a recently developed unified framework for instrument control. Following a MS1 survey scan (see below for LC-MS/MS details), the implemented acquisition workflow incorporated precursor deisotoping before the ML-based prediction, prioritization and selection of glycopeptide precursor ions for fragmentation. The glycopeptide classifier was employed only on precursors with intensities above 5,000 and charge states of 2–5 as these in our experience are more likely to generate high-quality interpretable MS/MS spectra. Only glycopeptide candidates with confidence scores exceeding the predefined threshold (for example ML0.95) were considered for MS/MS fragmentation. Dynamic exclusion of already fragmented candidates was applied for 8 s, preventing repeat fragmentation of the same precursor during the exclusion period, but typically allowing two or more isolations of an eluting analyte (for LC peaks with 10 s FWHM). Guided by the global and dataset-specific performance tests, different confidence score cut-offs were applied (ML0, ML0.15 and ML0.95). The on-line proof-of-concept experiments as well as the conventional intensity-based DDA experiments were repeated three times for each setting generating technical triplicates.

### Enrichment-free glycoproteomics of mouse plasma proteins

C57BL/6J mice (five males and five females, 8 weeks old) were purchased from Japan SLC (Shizuoka, Japan) and acclimatized for 2 weeks under controlled conditions (23 ± 2°C, 50 ± 10% relative humidity, 12 h light/dark cycle) with *ad libitum* access to food and water. At 10 weeks of age, mice were euthanized under deep isoflurane anesthesia. All animal procedures were approved by the Nagoya University Animal Care and Use Committee (A240095-003) and conducted in accordance with institutional and ARRIVE guidelines. Blood was collected from the right atrium into EDTA tubes immediately after euthanasia. Murine plasma was isolated by centrifugation (1,000 × g, 10 min, 4°C) and stored at −80°C. Protein concentration was determined using the BCA assay (Thermo Fisher Scientific). Plasma proteins (100 µg) were reduced using 10 mM DTT (30 min, 30°C), alkylated with 40 mM iodoacetamide (30 min, 20°C, in the dark), and the excess iodoacetamide was quenched with additional DTT. Proteins were digested overnight with sequencing-grade porcine trypsin (1:50, enzyme, w/w; Promega) at 37°C. Digestion was terminated by addition of TFA to 1% (v/v, final concentration), and peptides were desalted using Oligo R3 reversed-phase SPE microcolumns and dried. The plasma peptide mixtures were resuspended in water to a peptide concentration of 2 µg/µL. An aliquot was subsequently diluted to 0.25 µg/µL in 0.1% (v/v) formic acid for LC–MS/MS analysis.

Serving as a quantitative control, the plasma peptide samples from each of the five male and five female mice were separately labelled with tandem mass tags (TMT) according to the manufacturer’s protocol and as described^12^. Glycopeptides were enriched using custom-made hydrophilic interaction liquid chromatography (HILIC) SPE micro-columns packed with zwitterionic ZIC-HILIC resin (10 μm particle size, 200 Å pore size, kindly provided by Merck).

To predict murine plasma glycopeptide precursor ions during data acquisition of the unlabeled and unenriched peptide mixtures, model 1 was employed using the settings for the on-line ML-aided method described above. An ML threshold of 0.95 (ML0.95) was used to select high-confidence glycopeptides. Each plasma sample was analyzed in a single run generating five biological replicates of each sex. The TMT-labelled sample enriched for glycopeptides (n = 5/sex biological replicates) was analyzed without technical replicates using conventional intensity-based DDA on the Thermo Orbitrap Eclipse (see details below).

### LC-MS/MS settings

As mentioned above, different LC-MS/MS setups were used for the proof-of-concept and application experiments including 1) conventional intensity-based DDA performed on a Thermo Orbitrap Exploris (1a) or Thermo Orbitrap Eclipse (1b), and ML-aided DDA performed in 2) an off-line (static) configuration on a Thermo Orbitrap Exploris and 3) an on-line (real-time) configuration on a Thermo Orbitrap Eclipse both of which were applied to human sera (3a) whereas only on-line ML-aided DDA was applied to murine plasma (3b). To enable direct comparisons between the experimental conditions, the same amount of the same peptide sample was analyzed in technical (human sera) and biological (murine plasma) replicates across the different setups.

1. Conventional intensity-based DDA was performed using two different LC-MS/MS instruments i.e. Thermo Orbitrap Exploris 240 mass spectrometer (1a) or a Thermo Orbitrap Eclipse mass spectrometer (1b) to allow direct comparison to the off- and on-line ML-aided DDA methods acquired across these two instruments.

1b) For intensity-based DDA on the Orbitrap Exploris 240, peptides were loaded on a PepMap™ Neo Nano Trap Cartridge (5 mm × 300 μm inner diameter, Thermo) and separated on an C_18_ analytical column (Aurora Ultimate; 25 cm × 75 μm inner diameter, 1.7 μm particle size, IonOpticks) at 400 nL/min provided by a Vanquish Neo UHPLC System (Thermo). The binary mobile phase gradient was: 3-35% B over 90 min, 35-50% B over 8 min, 50-90% B over 2 min, and 10 min at 95% B. Solvent A: 0.1% (v/v) formic acid in water. Solvent B: 0.1% (v/v) formic acid in acetonitrile. The Orbitrap operating in positive ion polarity mode was used to acquire full MS1 scans (*m/z* 380-1,800, AGC: standard, 100 ms maximum accumulation time, 120,000 FWHM resolution at *m/z* 200). Employing data-dependent acquisition within a fixed 3 s cycle time, the N most abundant precursor ions from each MS full scan were prioritized to be isolated and fragmented utilizing higher-energy collision-induced dissociation (HCD, NCE 30%). Only multi-charged precursors (z = 2-7) were selected for fragmentation using an *m/z* 1.4 precursor ion isolation window. Fragment spectra were acquired in the Orbitrap (15,000 resolution, AGC: standard, maximum injection time mode “auto”, 20 s dynamic exclusion after a single isolation/fragmentation of a given precursor.

1b) For intensity-based DDA on the Orbitrap Eclipse, peptides were loaded on a PepMap™ Neo Nano Trap Cartridge (5 mm × 300 μm inner diameter, 5 μm particle, Thermo) and separated on an C_18_ analytical column (25 cm × 75 μm inner diameter, 1.9 μm particle, Nikkyo Technos) at 300 nL/min provided by an UltiMate 3000 RSLCnano System (Thermo). The binary mobile phase gradient was: 3% B over 10 min, 3-35% B over 90 min, 35-70% B over 3 min, 70% B over 3 min, 70-3% B over 4 min and 25 min at 3% B. Solvent A: 0.1% (v/v) formic acid in water. Solvent B: 0.1% (v/v) formic acid in acetonitrile. The Orbitrap Eclipse operating in positive ion polarity mode was used to acquire full MS1 scans (*m/z* 377-2,000, AGC: 100% (4 x 10^5^ ions), maximum injection time: 1,014 ms, resolution: 500,000). Employing data-dependent acquisition within a fixed 3 s cycle time, data-dependent acquisition of the TopN (N up to 20) most abundant precursor ions from each MS1 full scan were performed utilizing HCD with an NCE of 30%. Only multi-charged precursors (z = 2-5) were selected for fragmentation using an *m/z* 1.6 precursor ion isolation window. Fragment spectra were acquired in the Orbitrap (*m/z* 130-2,000, resolution: 3,000, AGC: 100% (5 x 10^4^ ions), maximum injection time: 100 ms, 8 s dynamic exclusion after a single isolation/fragmentation of a given precursor.

2) For the off-line proof-of-concept experiments of human sera, ML-aided DDA was carried out on a Thermo Orbitrap Exploris 240, which involved multiple steps: a) Peptide samples of human sera were analyzed by LC-MS runs exclusively collecting MS1 survey scans, b) Off-line interpretation of the MS1 data was performed by the glycopeptide classifier, and c) Glycopeptide candidates were predicted (ML0.95) and candidates compiled in an inclusion list embedded in a DDA method that was applied to a second injection of the same sample. Chromatographic conditions and mass spectrometer settings were identical to those described above for the conventional intensity-based DDA performed on the Orbitrap Exploris 240 with the following exceptions: Peptide samples were analyzed in MS1-only mode with data-dependent MS/MS acquisition disabled. Dynamic exclusion was excluded to maximize the fragmentation of precursors in the inclusion list using an Orbitrap resolution of 120,000.

3a) For the on-line proof-of-concept experiments of human sera, ML-aided DDA was carried out on a Thermo Orbitrap Eclipse. The Orbitrap Eclipse was used to acquire full MS1 scans (*m/z* 377-2,000, AGC: 100%, maximum injection time: 1,024 ms, resolution: 500,000). After acquisition of the MS1 survey scan, a deisotoping step was performed, and precursor ions were filtered to retain only multi-charged species (z = 2–5) with intensities exceeding 5,000. These precursor ions were then evaluated by the ML model. For the glycopeptide candidates with prediction scores above the specified threshold (i.e. ML0, ML0.15, or ML0.95), the Top-N most intense candidates were prioritized to be fragmented (N up to 15 for ML0 and ML0.95; N up to 20 for ML0.15) utilizing HCD with a NCE of 30% employing data-dependent acquisition within a fixed 3 s cycle time. Fragmentation was performed using an isolation window of 1.6 *m/z*. Fragment spectra were acquired in the Orbitrap (resolution = 30,000; AGC target = 50,000; maximum injection time of 54 ms for ML0.15 and maximum injection time of 100 ms for ML0 and ML0.95, with an 8 s dynamic exclusion applied after a single isolation/fragmentation of a given precursor.

3b) For the biological discovery of sex differences in murine plasma glycosylation, the on-line ML-aided DDA method was carried out using a Thermo Orbitrap Eclipse employing the same settings as described in 3a.

### Proof-of-concept for enrichment-free serum glycoproteomics

The proof-of-concept experiments performed on human sera were evaluated by 1) directly interrogating the prevalence of glycopeptide MS/MS spectra from the generated raw LC-MS/MS files and 2) analyzing the resulting glycopeptide-spectrum matches (glycoPSMs) and non-glycosylated PSMs after database searches.

For 1), the prevalence of glycopeptide MS/MS spectra was assessed using GlyCounter^20^ (v1.0.12). Raw files were processed with a signal-to-noise threshold of 3, and an intensity threshold of 1,000 for the fragment ions. HCD-MS/MS spectra were interrogated for diagnostic glycopeptide-specific oxonium ions, including HexNAc fragment ions (*m/z* 138.055 and *m/z* 204.087), sialic acid ions (*m/z* 274.092 and 292.103), and HexHexNAc ions (*m/z* 366.140). Spectra were classified as glycopeptide MS/MS spectra when at least three diagnostic oxonium ions were detected within a mass tolerance of 15 ppm in each product ion spectrum.

For 2), raw LC-MS/MS files were converted to mzXML format using RawConverter (v1.2.1) with a charge state range of 2–7 and the “Select monoisotopic *m/z* in DDA” option enabled. Glycopeptides and non-glycosylated peptides were identified directly from converted LC-MS/MS files using Byonic v5.8 (Protein Metrics). Searches were performed against the complete human reference proteome (*homo sapien*s, 20,431 entries, UniProtKB, March 2026) using a library of 288 N-glycan compositions for the off-line proof-of-concept experiments (**Supplementary Table S3**) and 200 N-glycan compositions for the on-line proof-of-concept experiments of human sera (**Supplementary Table S4**). Searches were conducted with standard precursor assignment settings or with the original precursor and charge assignments option enabled to preserve the precursor information reported by the instrument. Searches were directed against N-glycosylation (rare) allowing peptides to also carry Met oxidation (rare) as a variable modification. Each peptide was permitted to carry 1 rare and 2 common modifications. Cys alkylation was considered a fixed modification. N-ragged trypsin specificity was enabled to allow semi-specific cleavage at the N-terminus to improve the identification of peptides exhibiting non-tryptic N-terminal processing as is commonly observed when analyzing neat serum peptide mixtures. Confidence was achieved by consistently applying a PEP2D threshold of < 0.01 to all search outputs.

Potential bias was first assessed by comparing the observed glycoforms to the known N-glycome of human serum^10^. For this, the N-glycan compositions were extracted from the glycoPSMs identified from human serum and their relative frequencies were determined regardless of their protein carriers or glycosylation sites. This glycan distribution was compared to reported glycome profiles of human serum^10^. Secondly, the glycoproteins and non-glycosylated proteins identified in the proof-of-concept experiments were mapped on the abundance spectrum of known serum proteins^11^.

### Application of ML-aided DDA to murine plasma

The application experiments were evaluated by comparing ML-aided DDA with conventional DDA in two aspects: 1) the number and proportion of identified glycoPSMs and unique glycoproteins detected; and 2) the ability of the resulting data to reveal sex-related differences between plasma proteins from female and male mice. Leveraging the increased glycopeptide and glycoprotein coverage achieved by the ML-aided method, we further performed quantitative comparisons between female and male plasma to identify sex-associated differences in glycosylation, which were visualized using volcano plots and boxplots.

For 1), Glycopeptides were identified and quantified from the HCD-MS/MS data using Byonic v5.9.5 (Protein Metrics) operated as a node in Proteome Discoverer v.3.0. Search parameters included the use of i) the *Mus musculus* proteome database (17,150 reviewed sequences, UniProtKB, released June 2023), ii) an N-glycan search space comprising 288 predefined mammalian structures available in Byonic (excluding sodium adducts, **Supplementary Table S3**), iii) a search strategy permitting up to one N-glycan per peptide as a “rare” variable modification, fully-trypsin cleavage patterns with a maximum of two missed tryptic cleavages per peptide, up to 10/20 ppm mass tolerance of precursor/product ions from expected values, respectively, and up to one Met oxidation (+15.994 Da) per peptide (variable “common” modification), and by using monoisotopic correction (error check = +/- floor (mass in Da / 4,000)), and a decoy and default contaminant database available in Byonic. All searches were filtered to <1% false discovery rate (FDR) at the glycoprotein and glycopeptide level using a protein decoy database. Only confident glycopeptide identifications (PEP 2D < 0.01) were considered. Quantitation of (glyco)peptides was based on the intensity of precursor ion *m/z* as determined by the Minora Feature Detector with the minimum trace length set to 5 and maximum delta retention time of isotope pattern multiplets of 0.2 min within Proteome Discoverer v3.2.0.450. The feature mapper tool was enabled to allow chromatographic alignment of signals and extraction of precursor area based on a matched retention time and *m/z* within a time and mass tolerance window determined automatically by the software. Minimum S/N thresholds were enabled.

Abundances were converted to per-sample percentages, log2-transformed and centered on each sample’s mean (CLR). PCA was performed on the CLR transformed data, separately for the conventional and ML-aided DDA runs. Comparative analyses of the resulting glycoproteomics data were performed at two levels: at the glycan composition level (summing abundances across each glycan composition that was quantified across all samples) and at the glycopeptide level (comparing the glycopeptide level abundance only for glycopeptides quantified across all samples). We applied the limma R package to the CLR matrices, fitting ∼ sex with empirical Bayes moderated t-statistics and male − female as the contrast. Volcano plots show log2 fold change against −log10 *p* value, with significance called at Benjamini–Hochberg adjusted *p* < 0.05 and the most significant features labelled. Boxplots reusing these CLR values and limma statistics, with all 10 non-missing samples overlaid as points, top features of changed glycans and glycopeptides were shown. Further, we compared the global fucosylation levels between the ML-aided DDA runs of unenriched plasma peptide mixtures and quantitative controls acquired using TMT-DDA of enriched fractions of the same plasma samples. Within each platform, abundances were summed per glycan composition and fucosylation taken as the percentage of total glycan signal from fucosylated compositions; sex was tested per platform by two-tailed unpaired Welch t-tests.

### Statistics

Results from the conventional intensity-based DDA method were statistically compared to results from the ML-aided off- and on-line DDA methods using two-tailed unpaired Welch t-tests with *p* < 0.05 defining the threshold for significance. The significance levels and numbers of technical and biological replicates have been provided for each analysis. Bar graphs show the mean and error bars plot the standard deviation of data points. Boxplots show the mean and whiskers indicate spread of data. Individual data points are shown for all quantitative plots. Simple linear regression analyses were performed using GraphPad Prism version 10.2.0 (Dotmatics, San Diego, CA).

## Declarations

The authors declare no conflict of interest related to this work.

## Acknowledgements

We acknowledge financial support from the Human Glycome Atlas Project. We are grateful for project support from Prof Jun Hirabayashi and Mr Thomas Reilly.

## References

1 Varki, A. et al. Essentials of Glycobiology (2022). 10.1101/9781621824213

2 Kissel, T., Toes, R. E. M., Huizinga, T. W. J. & Wuhrer, M. Glycobiology of rheumatic diseases. Nat Rev Rheumatol 19, 28–43 (2023). 10.1038/s41584-022-00867-4

3 Kawahara, R. et al. Community evaluation of glycoproteomics informatics solutions reveals high-performance search strategies for serum glycopeptide analysis. Nat Methods 18, 1304–1316 (2021). 10.1038/s41592-021-01309-x

4 Chau, T. H., Chernykh, A., Kawahara, R. & Thaysen-Andersen, M. Critical considerations in N-glycoproteomics. Curr Opin Chem Biol 73, 102272 (2023). 10.1016/j.cbpa.2023.102272

5 Stavenhagen, K. et al. Quantitative mapping of glycoprotein micro-heterogeneity and macro-heterogeneity: an evaluation of mass spectrometry signal strengths using synthetic peptides and glycopeptides. J Mass Spectrom 48, 627–639 (2013). 10.1002/jms.3210

6 Tjondro, H. C. et al. Hyper-truncated Asn355- and Asn391-glycans modulate the activity of neutrophil granule myeloperoxidase. J Biol Chem 296, 100144 (2021). 10.1074/jbc.RA120.016342

7 Riley, N. M., Bertozzi, C. R. & Pitteri, S. J. A Pragmatic Guide to Enrichment Strategies for Mass Spectrometry-Based Glycoproteomics. Mol Cell Proteomics 20, 100029 (2021). 10.1074/mcp.R120.002277

8 Bagdonaite, I. et al. Glycoproteomics. Nature Reviews Methods Primers 2, 48 (2022). 10.1038/s43586-022-00128-4

9 Froehlich, J. W. et al. A classifier based on accurate mass measurements to aid large scale, unbiased glycoproteomics. Mol Cell Proteomics 12, 1017–1025 (2013). 10.1074/mcp.M112.025494

10 Clerc, F. et al. Human plasma protein N-glycosylation. Glycoconj J 33, 309–343 (2016). 10.1007/s10719-015-9626-2

11 Dey, K. K. et al. Deep undepleted human serum proteome profiling toward biomarker discovery for Alzheimer’s disease. Clin Proteomics 16, 16 (2019). 10.1186/s12014-019-9237-1

12 Kawahara, R. et al. Multi-omics definition of the sex-specific glycoproteome of murine tissues. bioRxiv, 2026.2003.2010.710926 (2026). 10.64898/2026.03.10.710926

13 Yang, R. B. et al. FUT8-Dependent Core Fucosylation: Essential for Platelet Function and a Target in Thrombosis. Arterioscler Thromb Vasc Biol 46, e324757 (2026). 10.1161/ATVBAHA.126.324757

14 Chau, T. H. et al. Serum AGP-1-Le(x) Glycoforms Report on Survivorship of Patients with Septic Shock Upon Admission to Intensive Care Unit. Mol Cell Proteomics 25, 101470 (2026). 10.1016/j.mcpro.2025.101470

15 Bienes, K. M. et al. Extracellular vesicles display distinct glycosignatures in high-grade serous ovarian carcinoma. BBA Adv 7, 100140 (2025). 10.1016/j.bbadva.2025.100140

16 Kawahara, R. et al. Glycoproteome remodeling and organelle-specific N-glycosylation accompany neutrophil granulopoiesis. Proc Natl Acad Sci U S A 120, e2303867120 (2023). 10.1073/pnas.2303867120

17 Houlahan, C. B. et al. Analysis of the Healthy Platelet Proteome Identifies a New Form of Domain-Specific O-Fucosylation. Mol Cell Proteomics 23, 100717 (2024). 10.1016/j.mcpro.2024.100717

18 Hulstaert, N. et al. ThermoRawFileParser: Modular, Scalable, and Cross-Platform RAW File Conversion. J Proteome Res 19, 537–542 (2020). 10.1021/acs.jproteome.9b00328

19 Hoopmann, M. R., McGann, C. D., Canterbury, J. D., von Haller, P. D. & Schweppe, D. K. Real-Time Instrument Control across Multiple Orbitrap Platforms through a Single Software Interface. J Proteome Res 24, 5277–5281 (2025). 10.1021/acs.jproteome.5c00269

20 Kothlow, K. et al. Extracting Informative Glycan-Specific Ions From Glycopeptide MS/MS Spectra With GlyCounter. Mol Cell Proteomics 24, 101085 (2025). 10.1016/j.mcpro.2025.101085

