## Supplementary File for "Enrichment-free glycoproteomics harnessing real-time mass defect-driven glycopeptide classification reveals sex differences in murine fucosylation"

**for**

Running title: Glycopeptide classifier facilitates enrichment-free glycoproteomics

Keywords: Enrichment, fucosylation, glycopeptide, glycoproteomics, machine learning, mass defect, real-time prediction, sex biology

Manuscript category: Brief communication

Corresponding author details:

Professor Morten Thaysen-Andersen

Institute for Glyco-core Research

Nagoya University - Nagoya – Japan

Phone: + 61 2 9850 7487

ORCID: 0000-0001-8327-6843

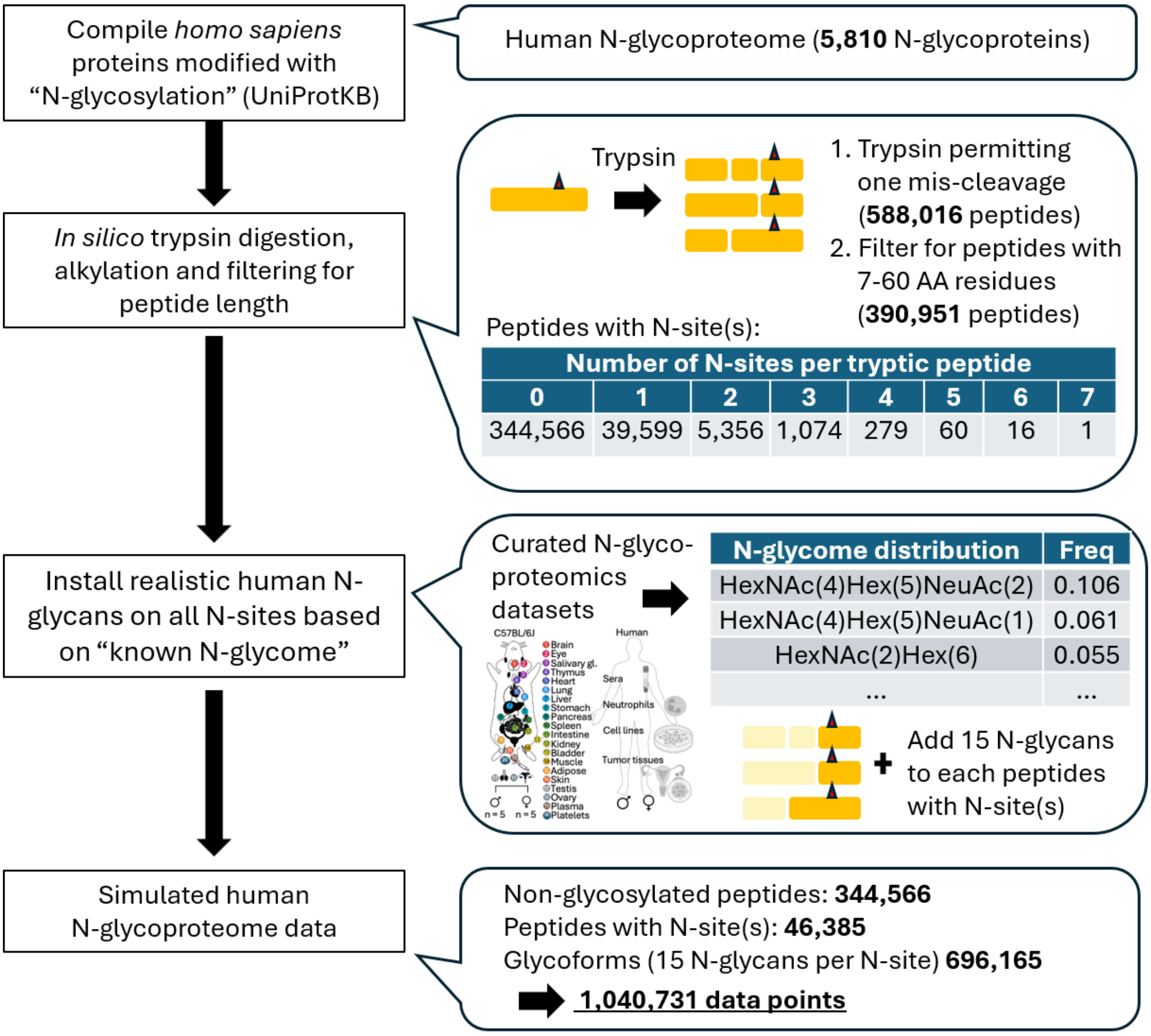


**Supplementary Figure S1**. **The simulated human N-glycoproteome**. Step-by-step overview of the *in silico* process of generating a simulated human N-glycoproteome used to train the glycopeptide classifier (model 2). See **Supplementary Table S1** for overview of curated N-glycoproteomics datasets used for the training of the experimental model (model 1).

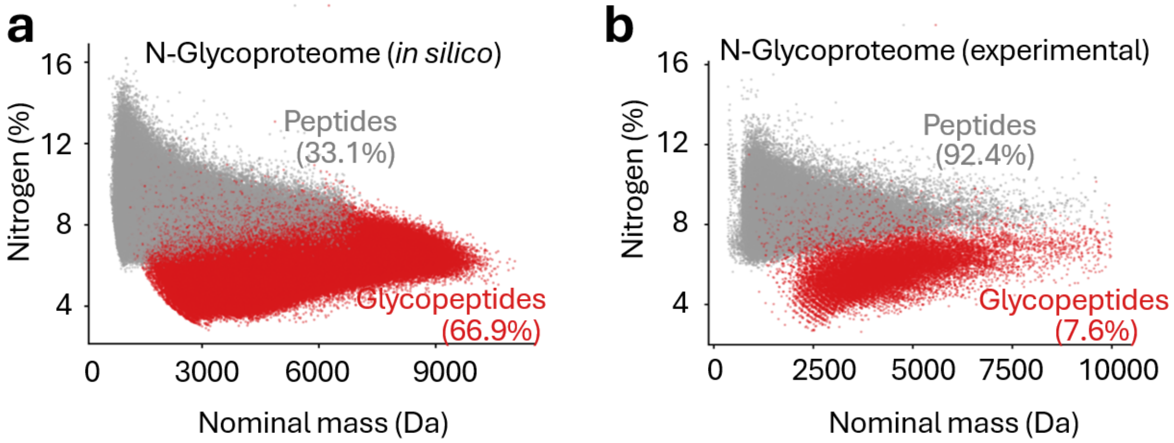


**Supplementary Figure S2**. **Tryptic N-glycopeptides exhibit suppressed nitrogen levels**. Global nitrogen-to-mass plots of tryptic N-glycopeptides (red) and non-glycosylated peptides (grey) from **a**) a simulated human N-glycoproteome (*in silico* generated, see **Supplementary Figure S1**) and **b**) experimental N-glycoproteomics data (see **Supplementary Table S1** for overview of curated N-glycoproteomics datasets used to train the experimental model). The relative proportion of data points labelled "glycopeptides" and non-glycosylated "peptides" are provided in brackets.

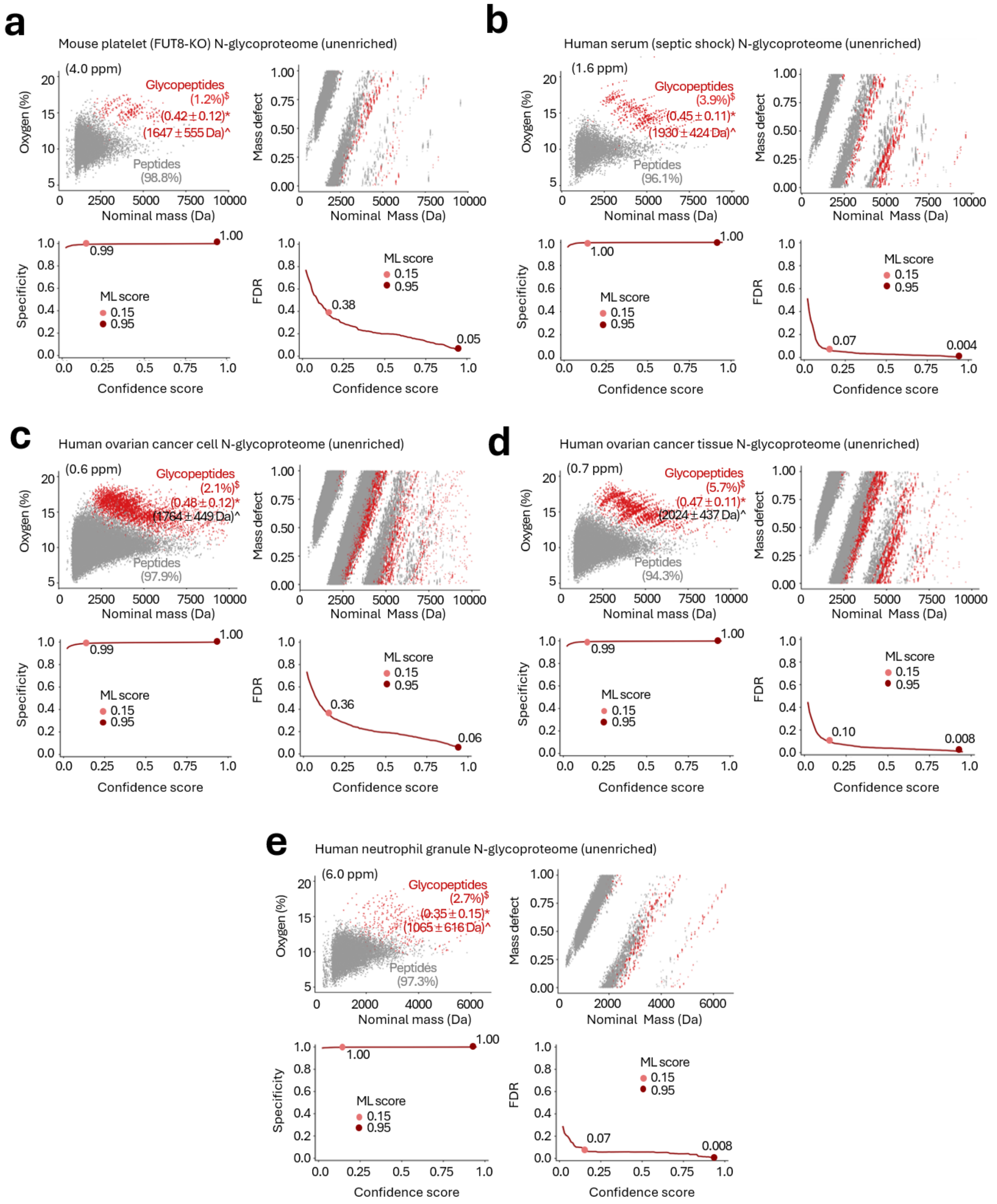


**Supplementary Figure S3**. **Dataset-specific performance of the glycopeptide classifier**. Oxygen levels, MD-to-nominal mass plots, prediction specificity and FDRs for N-glycoproteome data of unenriched peptide mixtures from **a**) mouse platelets (FUT8-KO), **b**) human serum (septic shock patients), **c**) human ovarian cancer cells, **d**) human ovarian tumor tissues, and **e**) human neutrophil granules. A fraction (70%) of data points from the selected datasets was used for the training of the glycopeptide classifier. Performance was tested on the remaining 30% of data points not previously seen by the ML model. In contrast, please note that the dataset-specific performance in **Figure 1h-i** was based on distinctly different datasets not used for training. [$]The relative proportion of data points are provided in brackets. *Glycan-to-molecular mass ratios (average ± SD) and ^glycan mass (average ± SD) of the identified N-glycopeptides.

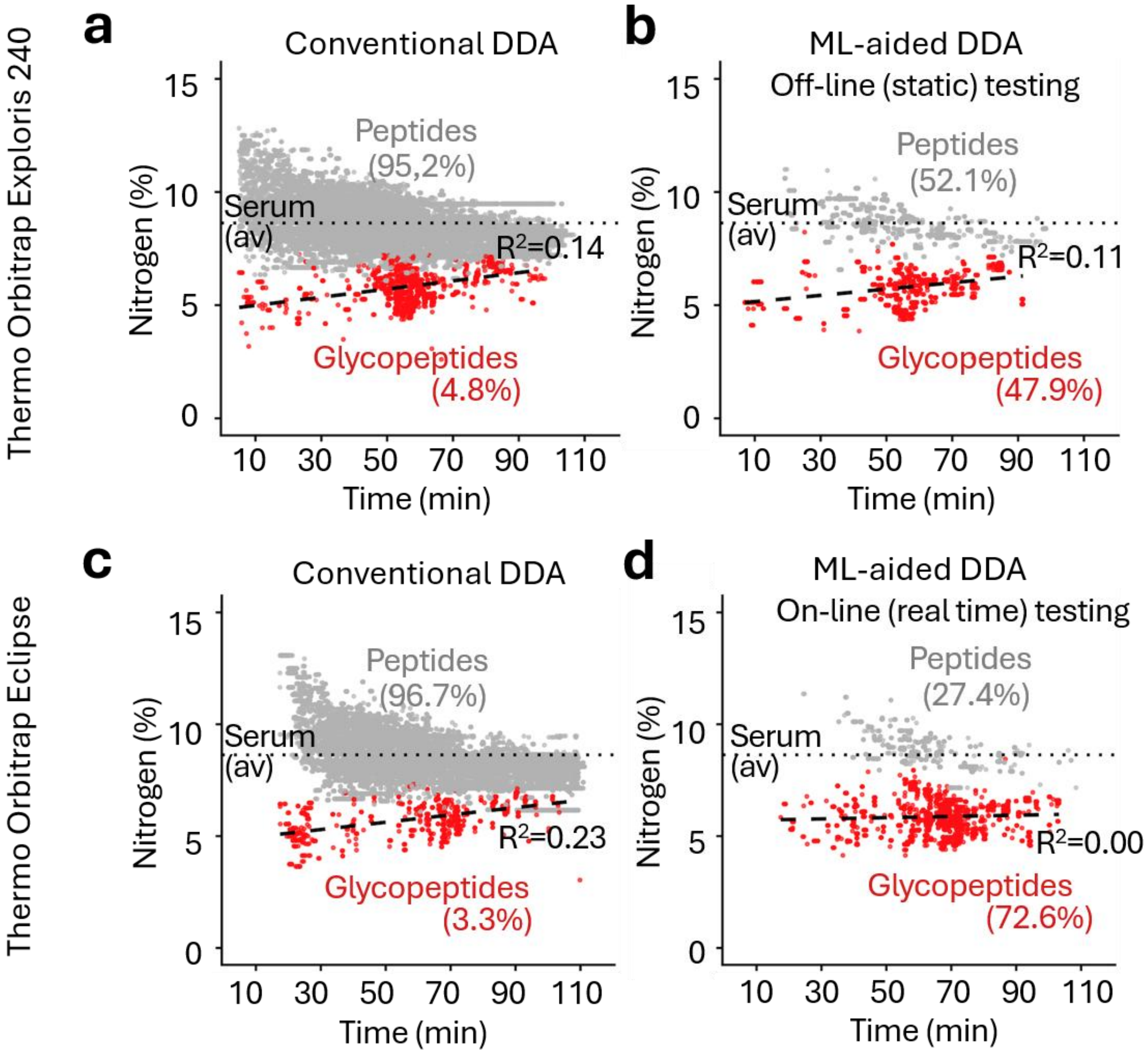


**Supplementary Figure S4. Suppressed nitrogen levels in the predicted N-glycopeptides**. Nitrogen-to-LC retention time plots of non-glycosylated peptides (grey) and N-glycopeptides (red) detected either with a Thermo Orbitrap Exploris 240 using **a)** conventional intensity-based DDA or **b**) off-line (static) ML-aided DDA or with a Thermo Orbitrap Eclipse using **c)** conventional intensity-based DDA or **d**) on-line (real-time) ML-aided DDA. Dotted grey line: average nitrogen content of tryptic peptides of human serum proteins[1]. The relative proportion of data points are provided in brackets. Black broken line: Trendline (linear regression, $R^2$) for glycopeptide data points.

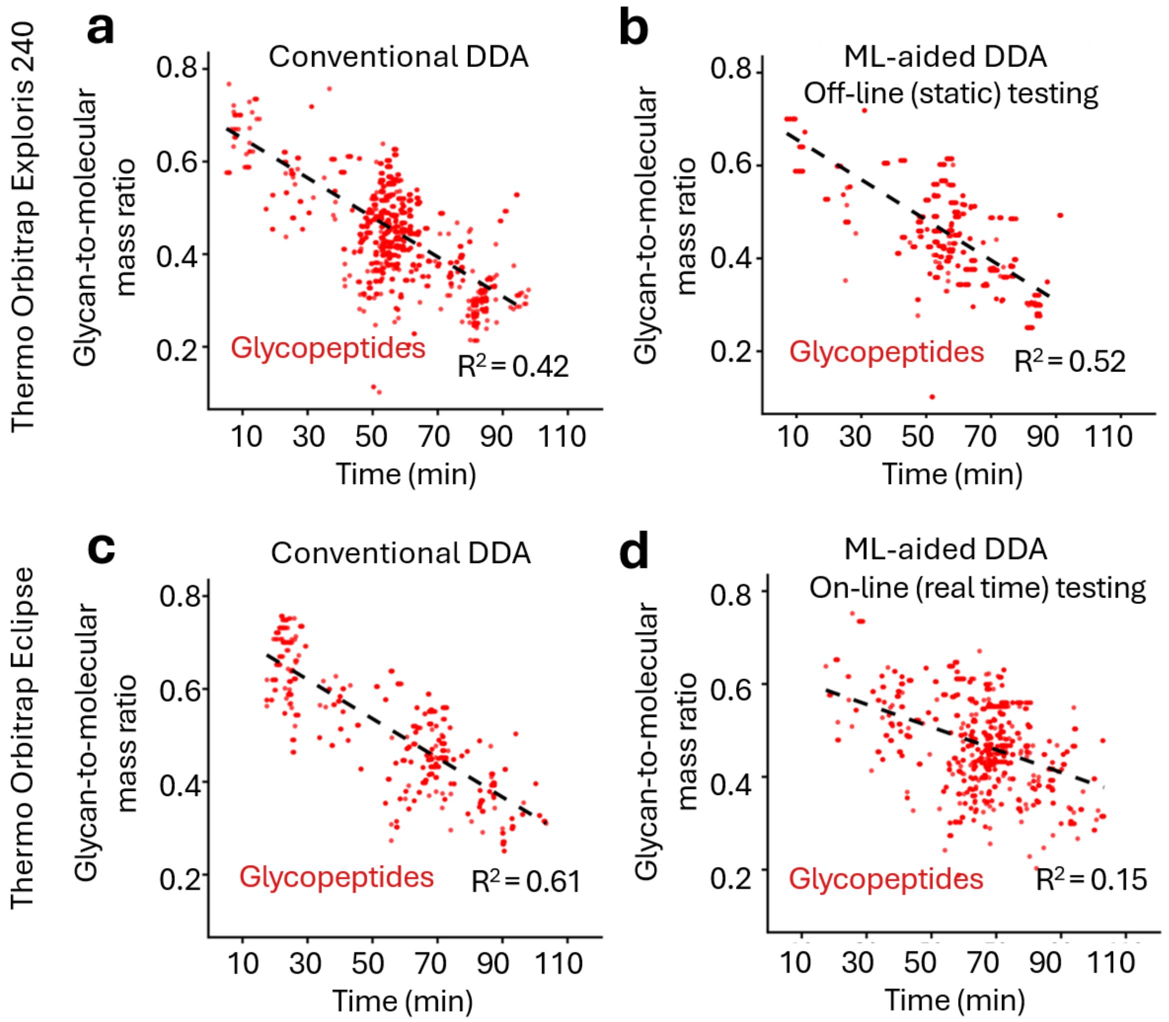


**Supplementary Figure S5**. **Glycan-to-molecular mass ratios differ over an LC-MS/MS run**. Glycan-to-molecular mass ratio as a function of LC retention time of N-glycopeptides detected either with a Thermo Orbitrap Exploris 240 using **a)** conventional intensity-based DDA or **b**) off-line (static) ML-aided DDA or with a Thermo Orbitrap Eclipse using **c)** conventional intensity-based DDA or **d**) on-line (real-time) ML-aided DDA. Black broken line: Trendline (linear regression, $R^2$) for glycopeptide data points.

**Supplementary Table S1**. Overview of the N-glycoproteomics datasets used for the training of the ML-driven glycopeptide classifier. *Average absolute values. ^Brain, eye, salivary gland, thymus, heart, lung, liver, stomach, pancreas, spleen, intestine, kidney, bladder, muscle, adipose, skin, testis, ovary, plasma[2]. [$] To avoid redundancy and biased training, the ~5.0m (glyco)PSMs were reduced to ~1.9m unique data points prior to the training.

| Tissues | Species | Condition(s) | Fraction | Non-glycoPSMs | GlycoPSMs | Total PSMs | Glycopeptide ratio | Δmass (ppm)* | Glycan mass (Da) | LC-MS/MS files |
|---|---|---|---|---|---|---|---|---|---|---|
| Wide tissue panel^ | Mouse | C57BL/6, normal | Enriched | 51,230 | 34,321 | 85,551 | 40.12% | 4.04 | 1,789 | 20 |
| Platelets | Mouse | FUT8-KO, wildtype | Enriched | 61,541 | 9,761 | 71,302 | 13.69% | 1.71 | 1,635 | 6 |
| Platelets | Mouse | FUT8-KO, wildtype | Flow through | 153,431 | 189 | 153,620 | 0.12% | 1.86 | 774 | 6 |
| Platelets | Mouse | FUT8-KO, wildtype | Non-enriched | 151,453 | 1,419 | 152,872 | 0.93% | 1.57 | 1,626 | 6 |
| Sera | Human | Septic shock, healthy | Enriched | 216,521 | 118,094 | 334,615 | 35.29% | 1.07 | 2,131 | 37 |
| Sera | Human | Septic shock, healthy | Non-enriched | 573,510 | 9,458 | 582,968 | 1.62% | 1.04 | 1,907 | 37 |
| Ovarian cell lines | Human | Ovarian cancer | Non-enriched | 2,466,453 | 27,072 | 2,493,525 | 1.09% | 0.57 | 1,745 | 60 |
| Ovarian tissues | Human | Ovarian cancer | Non-enriched | 839,217 | 32,915 | 872,132 | 3.77% | 0.66 | 1,982 | 32 |
| Neutrophil granules | Human | Healthy donor | Enriched | 40,949 | 14,545 | 55,494 | 26.21% | 1.31 | 1,203 | 15 |
| Neutrophil granules | Human | Healthy donor | Non-enriched | 158,632 | 3,106 | 161,738 | 1.92% | 6.04 | 1,043 | 15 |
| | | | **Total** | **4,712,937** | **250,880** | **4,96,3817**[$] | **5.05%** | **1.99** | **1,584** | **234** |

**Supplementary Table S2**. Overview of N-glycoproteomics datasets previous unseen by the ML-driven glycopeptide classifier used for independent testing. *Average absolute values.

| Tissues | Species | Condition(s) | Fraction | Non-glycoPSMs | GlycoPSMs | Total PSMs | Glycopeptide ratio | Δmass (ppm)* | Glycan mass (Da) | LC-MS/MS files |
|---|---|---|---|---|---|---|---|---|---|---|
| Platelets | Human | Healthy donor | Unenriched | 274,962 | 3,747 | 278,709 | 1.34% | 1.61 | 1,945 | 22 |
| Serum | Human | Ovarian cancer | Unenriched | 24,535 | 1,876 | 26,411 | 7.10% | 1.02 | 2,069 | 3 |
| | | | **Total** | **299,497** | **5,623** | **305,120** | **1.84** | **1.56** | **1,986** | **25** |

**Supplementary Table S3**. Glycan search space (288 N-glycan compositions) used for the Byonic searches of mouse tissue samples for the off-line proof-of-concept experiments. Masses are provided as neutral monoisotopic values (M).

| Glycan composition | Mass (Da) |
|---|---|
| HexNAc(1) | 203.0794 |
| HexNAc(2) | 406.1587 |
| HexNAc(2)Hex(1) | 568.2116 |
| HexNAc(2)Hex(2) | 730.2644 |
| HexNAc(1)Fuc(1) | 349.1373 |
| HexNAc(2)Fuc(1) | 552.2167 |
| HexNAc(2)Hex(1)Fuc(1) | 714.2695 |
| HexNAc(2)Hex(2)Fuc(1) | 876.3223 |
| HexNAc(2)Hex(3)Fuc(1) | 1038.3750 |
| HexNAc(2)Hex(3) | 892.3172 |
| HexNAc(2)Hex(4) | 1054.3700 |
| HexNAc(3)Hex(3) | 1095.3966 |
| HexNAc(2)Hex(3)Fuc(2) | 1184.4330 |
| HexNAc(2)Hex(4)Fuc(1) | 1200.4280 |
| HexNAc(2)Hex(5) | 1216.4229 |
| HexNAc(3)Hex(3)Fuc(1) | 1241.4545 |
| HexNAc(3)Hex(4) | 1257.4494 |
| HexNAc(4)Hex(3) | 1298.4760 |
| HexNAc(2)Hex(5)Fuc(1) | 1362.4808 |
| HexNAc(2)Hex(6) | 1378.4757 |
| HexNAc(3)Hex(3)Fuc(2) | 1387.5124 |
| HexNAc(3)Hex(4)Fuc(1) | 1403.5073 |
| HexNAc(3)Hex(5) | 1419.5022 |
| HexNAc(4)Hex(3)Fuc(1) | 1444.5339 |
| HexNAc(4)Hex(4) | 1460.5288 |
| HexNAc(5)Hex(3) | 1501.5553 |
| HexNAc(2)Hex(7) | 1540.5285 |
| HexNAc(3)Hex(4)NeuAc(1) | 1548.5448 |
| HexNAc(3)Hex(4)Fuc(2) | 1549.5652 |
| HexNAc(3)Hex(4)NeuGc(1) | 1564.5397 |
| HexNAc(3)Hex(5)Fuc(1) | 1565.5601 |
| HexNAc(3)Hex(6) | 1581.5551 |
| HexNAc(4)Hex(3)NeuAc(1) | 1589.5714 |
| HexNAc(4)Hex(3)Fuc(2) | 1590.5918 |
| HexNAc(4)Hex(3)NeuGc(1) | 1605.5663 |
| HexNAc(4)Hex(4)Fuc(1) | 1606.5867 |
| HexNAc(4)Hex(5) | 1622.5816 |
| HexNAc(5)Hex(3)Fuc(1) | 1647.6132 |
| HexNAc(5)Hex(4) | 1663.6082 |
| HexNAc(3)Hex(4)Fuc(1)NeuAc(1) | 1694.6027 |
| HexNAc(2)Hex(8) | 1702.5813 |
| HexNAc(6)Hex(3) | 1704.6347 |
| HexNAc(3)Hex(4)Fuc(1)NeuGc(1) | 1710.5977 |
| HexNAc(3)Hex(5)NeuAc(1) | 1710.5977 |
| HexNAc(3)Hex(5)NeuGc(1) | 1726.5926 |
| HexNAc(3)Hex(6)Fuc(1) | 1727.6130 |
| HexNAc(4)Hex(3)Fuc(3) | 1736.6497 |
| HexNAc(4)Hex(4)NeuAc(1) | 1751.6242 |
| HexNAc(4)Hex(4)Fuc(2) | 1752.6446 |
| HexNAc(4)Hex(4)NeuGc(1) | 1767.6191 |
| HexNAc(4)Hex(5)Fuc(1) | 1768.6395 |
| HexNAc(4)Hex(6) | 1784.6344 |
| HexNAc(5)Hex(4)Fuc(1) | 1809.6661 |
| HexNAc(5)Hex(5) | 1825.6610 |
| HexNAc(6)Hex(3)Fuc(1) | 1850.6926 |
| HexNAc(2)Hex(9) | 1864.6342 |
| HexNAc(6)Hex(4) | 1866.6875 |
| HexNAc(3)Hex(6)NeuAc(1) | 1872.6505 |
| HexNAc(3)Hex(6)NeuGc(1) | 1888.6454 |
| HexNAc(4)Hex(4)Fuc(1)NeuAc(1) | 1897.6821 |
| HexNAc(7)Hex(3) | 1907.7141 |
| HexNAc(4)Hex(4)Fuc(1)NeuGc(1) | 1913.6770 |
| HexNAc(4)Hex(5)NeuAc(1) | 1913.6770 |
| HexNAc(4)Hex(5)Fuc(2) | 1914.6974 |
| HexNAc(4)Hex(5)NeuGc(1) | 1929.6719 |
| HexNAc(4)Hex(6)Fuc(1) | 1930.6923 |
| HexNAc(5)Hex(3)Fuc(1)NeuAc(1) | 1938.7087 |
| HexNAc(4)Hex(7) | 1946.6873 |
| HexNAc(5)Hex(4)NeuAc(1) | 1954.7036 |
| HexNAc(5)Hex(3)Fuc(1)NeuGc(1) | 1954.7036 |
| HexNAc(5)Hex(4)Fuc(2) | 1955.7240 |
| HexNAc(5)Hex(4)NeuGc(1) | 1970.6985 |
| HexNAc(5)Hex(5)Fuc(1) | 1971.7189 |
| HexNAc(5)Hex(6) | 1987.7138 |
| HexNAc(6)Hex(3)Fuc(2) | 1996.7505 |
| HexNAc(6)Hex(4)Fuc(1) | 2012.7454 |
| HexNAc(3)Hex(6)Fuc(1)NeuAc(1) | 2018.7084 |

| | |
|---|---|
| HexNAc(2)Hex(10) | 2026.6870 |
| HexNAc(6)Hex(5) | 2028.7404 |
| HexNAc(3)Hex(6)Fuc(1)NeuGc(1) | 2034.7033 |
| HexNAc(4)Hex(4)Fuc(2)NeuAc(1) | 2043.7400 |
| HexNAc(7)Hex(3)Fuc(1) | 2053.7720 |
| HexNAc(4)Hex(5)Fuc(1)NeuAc(1) | 2059.7349 |
| HexNAc(4)Hex(4)Fuc(2)NeuGc(1) | 2059.7349 |
| HexNAc(4)Hex(5)Fuc(3) | 2060.7553 |
| HexNAc(7)Hex(4) | 2069.7669 |
| HexNAc(4)Hex(6)NeuAc(1) | 2075.7299 |
| HexNAc(4)Hex(5)Fuc(1)NeuGc(1) | 2075.7299 |
| HexNAc(4)Hex(6)Fuc(2) | 2076.7503 |
| HexNAc(4)Hex(6)NeuGc(1) | 2091.7248 |
| HexNAc(4)Hex(7)Fuc(1) | 2092.7452 |
| HexNAc(5)Hex(4)Fuc(1)NeuAc(1) | 2100.7615 |
| HexNAc(8)Hex(3) | 2110.7935 |
| HexNAc(5)Hex(4)Fuc(1)NeuGc(1) | 2116.7564 |
| HexNAc(5)Hex(5)NeuAc(1) | 2116.7564 |
| HexNAc(5)Hex(5)Fuc(2) | 2117.7768 |
| HexNAc(5)Hex(5)NeuGc(1) | 2132.7513 |
| HexNAc(5)Hex(6)Fuc(1) | 2133.7717 |
| HexNAc(6)Hex(3)Fuc(1)NeuAc(1) | 2141.7880 |
| HexNAc(6)Hex(3)Fuc(3) | 2142.8084 |
| HexNAc(5)Hex(7) | 2149.7666 |
| HexNAc(6)Hex(4)NeuAc(1) | 2157.7830 |
| HexNAc(6)Hex(3)Fuc(1)NeuGc(1) | 2157.7830 |
| HexNAc(6)Hex(4)Fuc(2) | 2158.8034 |
| HexNAc(6)Hex(5)Fuc(1) | 2174.7983 |
| HexNAc(2)Hex(11) | 2188.7398 |
| HexNAc(6)Hex(6) | 2190.7932 |
| HexNAc(4)Hex(5)NeuAc(2) | 2204.7724 |
| HexNAc(4)Hex(5)Fuc(2)NeuAc(1) | 2205.7928 |
| HexNAc(4)Hex(5)Fuc(4) | 2206.8132 |
| HexNAc(7)Hex(4)Fuc(1) | 2215.8248 |
| HexNAc(4)Hex(5)NeuAc(1)NeuGc(1) | 2220.7674 |
| HexNAc(4)Hex(5)Fuc(2)NeuGc(1) | 2221.7878 |
| HexNAc(4)Hex(6)Fuc(1)NeuAc(1) | 2221.7878 |
| HexNAc(4)Hex(6)Fuc(3) | 2222.8082 |
| HexNAc(4)Hex(5)NeuGc(2) | 2236.7623 |
| HexNAc(4)Hex(6)Fuc(1)NeuGc(1) | 2237.7827 |
| HexNAc(4)Hex(7)NeuAc(1) | 2237.7827 |
| HexNAc(4)Hex(7)Fuc(2) | 2238.8031 |
| HexNAc(5)Hex(4)NeuAc(2) | 2245.7990 |
| HexNAc(5)Hex(4)Fuc(2)NeuAc(1) | 2246.8194 |

| | |
|---|---|
| HexNAc(8)Hex(3)Fuc(1) | 2256.8514 |
| HexNAc(5)Hex(4)NeuAc(1)NeuGc(1) | 2261.7939 |
| HexNAc(5)Hex(5)Fuc(1)NeuAc(1) | 2262.8143 |
| HexNAc(5)Hex(4)Fuc(2)NeuGc(1) | 2262.8143 |
| HexNAc(5)Hex(5)Fuc(3) | 2263.8347 |
| HexNAc(8)Hex(4) | 2272.8463 |
| HexNAc(5)Hex(4)NeuGc(2) | 2277.7888 |
| HexNAc(5)Hex(5)Fuc(1)NeuGc(1) | 2278.8092 |
| HexNAc(5)Hex(6)NeuAc(1) | 2278.8092 |
| HexNAc(5)Hex(6)Fuc(2) | 2279.8296 |
| HexNAc(6)Hex(3)Fuc(2)NeuAc(1) | 2287.8459 |
| HexNAc(5)Hex(6)NeuGc(1) | 2294.8041 |
| HexNAc(5)Hex(7)Fuc(1) | 2295.8245 |
| HexNAc(6)Hex(3)Fuc(2)NeuGc(1) | 2303.8409 |
| HexNAc(5)Hex(8) | 2311.8195 |
| HexNAc(9)Hex(3) | 2313.8728 |
| HexNAc(6)Hex(5)Fuc(2) | 2320.8562 |
| HexNAc(6)Hex(6)Fuc(1) | 2336.8511 |
| HexNAc(2)Hex(12) | 2350.7926 |
| HexNAc(4)Hex(5)Fuc(1)NeuAc(2) | 2350.8304 |
| HexNAc(6)Hex(7) | 2352.8460 |
| HexNAc(7)Hex(4)Fuc(2) | 2361.8827 |
| HexNAc(4)Hex(5)Fuc(1)NeuAc(1)NeuGc(1) | 2366.8253 |
| HexNAc(4)Hex(5)Fuc(1)NeuGc(2) | 2382.8202 |
| HexNAc(5)Hex(4)Fuc(1)NeuAc(2) | 2391.8569 |
| HexNAc(7)Hex(6) | 2393.8726 |
| HexNAc(5)Hex(5)NeuAc(2) | 2407.8518 |
| HexNAc(5)Hex(4)Fuc(1)NeuAc(1)NeuGc(1) | 2407.8518 |
| HexNAc(5)Hex(4)Fuc(1)NeuGc(2) | 2423.8467 |
| HexNAc(5)Hex(5)NeuAc(1)NeuGc(1) | 2423.8467 |
| HexNAc(5)Hex(6)Fuc(1)NeuAc(1) | 2424.8671 |
| HexNAc(5)Hex(6)Fuc(3) | 2425.8875 |
| HexNAc(6)Hex(3)Fuc(1)NeuAc(2) | 2432.8835 |
| HexNAc(8)Hex(5) | 2434.8991 |
| HexNAc(5)Hex(5)NeuGc(2) | 2439.8416 |
| HexNAc(5)Hex(6)Fuc(1)NeuGc(1) | 2440.8620 |
| HexNAc(6)Hex(3)Fuc(1)NeuAc(1)NeuGc(1) | 2448.8784 |
| HexNAc(5)Hex(8)Fuc(1) | 2457.8774 |
| HexNAc(9)Hex(3)Fuc(1) | 2459.9307 |
| HexNAc(6)Hex(3)Fuc(1)NeuGc(2) | 2464.8733 |
| HexNAc(9)Hex(4) | 2475.9257 |
| HexNAc(6)Hex(6)Fuc(2) | 2482.9090 |
| HexNAc(6)Hex(7)Fuc(1) | 2498.9039 |

| | |
|---|---|
| HexNAc(7)Hex(6)Fuc(1) | 2539.9305 |
| HexNAc(5)Hex(5)Fuc(1)NeuAc(2) | 2553.9097 |
| HexNAc(7)Hex(7) | 2555.9254 |
| HexNAc(5)Hex(5)Fuc(1)NeuAc(1)NeuGc(1) | 2569.9046 |
| HexNAc(5)Hex(6)NeuAc(2) | 2569.9046 |
| HexNAc(5)Hex(6)Fuc(2)NeuAc(1) | 2570.9250 |
| HexNAc(8)Hex(5)Fuc(1) | 2580.9570 |
| HexNAc(5)Hex(5)Fuc(1)NeuGc(2) | 2585.8996 |
| HexNAc(5)Hex(6)NeuAc(1)NeuGc(1) | 2585.8996 |
| HexNAc(5)Hex(7)Fuc(1)NeuAc(1) | 2586.9200 |
| HexNAc(5)Hex(6)Fuc(2)NeuGc(1) | 2586.9200 |
| HexNAc(8)Hex(6) | 2596.9519 |
| HexNAc(5)Hex(6)NeuGc(2) | 2601.8945 |
| HexNAc(5)Hex(9)Fuc(1) | 2619.9302 |
| HexNAc(9)Hex(4)Fuc(1) | 2621.9836 |
| HexNAc(6)Hex(6)Fuc(3) | 2628.9669 |
| HexNAc(4)Hex(5)Fuc(3)NeuAc(2) | 2642.9462 |
| HexNAc(4)Hex(5)Fuc(3)NeuAc(1) | 2351.8508 |
| HexNAc(6)Hex(7)NeuAc(1) | 2643.9414 |
| HexNAc(4)Hex(5)Fuc(3)NeuAc(1)NeuGc(1) | 2658.9411 |
| HexNAc(6)Hex(7)NeuGc(1) | 2659.9363 |
| HexNAc(4)Hex(5)Fuc(3)NeuGc(2) | 2674.9360 |
| HexNAc(6)Hex(9) | 2676.9517 |
| HexNAc(7)Hex(7)Fuc(1) | 2701.9833 |
| HexNAc(5)Hex(6)Fuc(1)NeuAc(2) | 2715.9626 |
| HexNAc(5)Hex(6)Fuc(3)NeuAc(1) | 2716.9830 |
| HexNAc(7)Hex(8) | 2717.9782 |
| HexNAc(5)Hex(6)Fuc(1)NeuAc(1)NeuGc(1) | 2731.9575 |
| HexNAc(5)Hex(6)Fuc(3)NeuGc(1) | 2732.9779 |
| HexNAc(5)Hex(6)Fuc(1)NeuGc(2) | 2747.9524 |
| HexNAc(8)Hex(7) | 2759.0048 |
| HexNAc(6)Hex(6)NeuAc(1) | 2481.8886 |
| HexNAc(6)Hex(6)NeuAc(2) | 2772.9840 |
| HexNAc(6)Hex(6)NeuAc(1)NeuGc(1) | 2788.9789 |
| HexNAc(6)Hex(7)Fuc(1)NeuAc(1) | 2789.9993 |
| HexNAc(9)Hex(6) | 2800.0313 |
| HexNAc(6)Hex(6)NeuGc(2) | 2804.9738 |
| HexNAc(6)Hex(7)Fuc(1)NeuGc(1) | 2805.9942 |
| HexNAc(6)Hex(8)NeuAc(1) | 2805.9942 |
| HexNAc(5)Hex(6)NeuAc(3) | 2861.0001 |
| HexNAc(7)Hex(8)Fuc(1) | 2864.0361 |
| HexNAc(5)Hex(6)NeuAc(2)NeuGc(1) | 2876.9950 |
| HexNAc(5)Hex(7)Fuc(1)NeuAc(2) | 2878.0154 |

| | |
|---|---|
| HexNAc(5)Hex(6)NeuAc(1)NeuGc(2) | 2892.9899 |
| HexNAc(5)Hex(7)Fuc(1)NeuAc(1)NeuGc(1) | 2894.0103 |
| HexNAc(5)Hex(8)Fuc(4) | 2896.0511 |
| HexNAc(5)Hex(6)NeuGc(3) | 2908.9848 |
| HexNAc(5)Hex(7)Fuc(1)NeuGc(2) | 2910.0052 |
| HexNAc(6)Hex(6)Fuc(1)NeuAc(2) | 2919.0419 |
| HexNAc(6)Hex(6)Fuc(1)NeuAc(1) | 2627.9465 |
| HexNAc(8)Hex(8) | 2921.0576 |
| HexNAc(6)Hex(6)Fuc(1)NeuAc(1)NeuGc(1) | 2935.0368 |
| HexNAc(6)Hex(7)NeuAc(2) | 2935.0368 |
| HexNAc(6)Hex(7)Fuc(4) | 2937.0776 |
| HexNAc(9)Hex(6)Fuc(1) | 2946.0892 |
| HexNAc(6)Hex(7)NeuAc(1)NeuGc(1) | 2951.0318 |
| HexNAc(6)Hex(6)Fuc(1)NeuGc(2) | 2951.0318 |
| HexNAc(6)Hex(8)Fuc(1)NeuAc(1) | 2952.0522 |
| HexNAc(6)Hex(7)NeuGc(2) | 2967.0267 |
| HexNAc(5)Hex(6)Fuc(1)NeuAc(3) | 3007.0580 |
| HexNAc(7)Hex(8)NeuAc(1) | 3009.0736 |
| HexNAc(5)Hex(6)Fuc(1)NeuAc(2)NeuGc(1) | 3023.0529 |
| HexNAc(7)Hex(8)NeuGc(1) | 3025.0685 |
| HexNAc(5)Hex(6)Fuc(1)NeuAc(1)NeuGc(2) | 3039.0478 |
| HexNAc(6)Hex(5)Fuc(1)NeuAc(2) | 2756.9891 |
| HexNAc(6)Hex(5)Fuc(1)NeuAc(1) | 2465.8937 |
| HexNAc(6)Hex(5)Fuc(1)NeuAc(3) | 3048.0845 |
| HexNAc(5)Hex(6)Fuc(1)NeuGc(3) | 3055.0427 |
| HexNAc(6)Hex(5)Fuc(1)NeuAc(1)NeuGc(1) | 2772.9840 |
| HexNAc(6)Hex(5)Fuc(1)NeuAc(2)NeuGc(1) | 3064.0794 |
| HexNAc(8)Hex(8)Fuc(1) | 3067.1155 |
| HexNAc(6)Hex(5)Fuc(1)NeuAc(1)NeuGc(2) | 3080.0743 |
| HexNAc(6)Hex(7)Fuc(1)NeuAc(2) | 3081.0947 |
| HexNAc(8)Hex(9) | 3083.1104 |
| HexNAc(6)Hex(7)Fuc(5) | 3083.1356 |
| HexNAc(6)Hex(5)Fuc(1)NeuGc(3) | 3096.0693 |
| HexNAc(6)Hex(11)Fuc(1) | 3147.1152 |
| HexNAc(10)Hex(7) | 3165.1635 |
| HexNAc(6)Hex(6)Fuc(1)NeuAc(3) | 3210.1373 |
| HexNAc(6)Hex(6)Fuc(1)NeuAc(2)NeuGc(1) | 3226.1323 |
| HexNAc(6)Hex(7)NeuAc(3) | 3226.1323 |
| HexNAc(6)Hex(7)Fuc(4)NeuAc(1) | 3228.1731 |

| | |
|---|---|
| HexNAc(8)Hex(9)Fuc(1) | 3229.1683 |
| HexNAc(6)Hex(7)NeuAc(2)NeuGc(1) | 3242.1272 |
| HexNAc(6)Hex(6)Fuc(1)NeuAc(1)NeuGc(2) | 3242.1272 |
| HexNAc(6)Hex(7)Fuc(4)NeuGc(1) | 3244.1680 |
| HexNAc(6)Hex(6)Fuc(1)NeuGc(3) | 3258.1221 |
| HexNAc(6)Hex(10)Fuc(1)NeuAc(1) | 3276.1578 |
| HexNAc(7)Hex(7)Fuc(1)NeuAc(2) | 3284.1741 |
| HexNAc(6)Hex(10)Fuc(1)NeuGc(1) | 3292.1527 |
| HexNAc(6)Hex(7)Fuc(1)NeuAc(3) | 3372.1902 |
| HexNAc(6)Hex(9)Fuc(1)NeuAc(2) | 3405.2004 |
| HexNAc(6)Hex(9)Fuc(1)NeuAc(1)NeuGc(1) | 3421.1953 |
| HexNAc(9)Hex(9)Fuc(1) | 3432.2477 |
| HexNAc(7)Hex(8)Fuc(1)NeuAc(1) | 3155.1315 |
| HexNAc(7)Hex(8)Fuc(1)NeuAc(2) | 3446.2269 |
| HexNAc(9)Hex(10) | 3448.2426 |
| HexNAc(7)Hex(8)Fuc(1)NeuAc(1)NeuGc(1) | 3462.2219 |
| HexNAc(7)Hex(8)Fuc(1)NeuGc(2) | 3478.2168 |
| HexNAc(6)Hex(7)NeuAc(4) | 3517.2277 |
| HexNAc(6)Hex(7)NeuAc(3)NeuGc(1) | 3533.2226 |
| HexNAc(6)Hex(7)NeuAc(2)NeuGc(2) | 3549.2175 |
| HexNAc(7)Hex(7)Fuc(1)NeuAc(3) | 3575.2695 |
| HexNAc(7)Hex(7)Fuc(1)NeuAc(2)NeuGc(1) | 3591.2645 |

| | |
|---|---|
| HexNAc(9)Hex(10)Fuc(1) | 3594.3005 |
| HexNAc(7)Hex(7)Fuc(1)NeuAc(1)NeuGc(2) | 3607.2594 |
| HexNAc(6)Hex(7)Fuc(1)NeuAc(4) | 3663.2856 |
| HexNAc(6)Hex(7)Fuc(1)NeuAc(3)NeuGc(1) | 3679.2805 |
| HexNAc(7)Hex(6)Fuc(1)NeuAc(4) | 3704.3121 |
| HexNAc(7)Hex(6)Fuc(1)NeuAc(3)NeuGc(1) | 3720.3070 |
| HexNAc(7)Hex(8)Fuc(1)NeuAc(3) | 3737.3224 |
| HexNAc(7)Hex(8)Fuc(1)NeuAc(2)NeuGc(1) | 3753.3173 |
| HexNAc(7)Hex(8)Fuc(1)NeuAc(1)NeuGc(2) | 3769.3122 |
| HexNAc(7)Hex(8)Fuc(1)NeuGc(3) | 3785.3071 |
| HexNAc(10)Hex(10)Fuc(1) | 3797.3799 |
| HexNAc(7)Hex(7)Fuc(1)NeuAc(4) | 3866.3650 |
| HexNAc(7)Hex(8)Fuc(1)NeuAc(4) | 4028.4178 |
| HexNAc(8)Hex(9)Fuc(1)NeuAc(3) | 4102.4546 |
| HexNAc(8)Hex(9)Fuc(1)NeuAc(2)NeuGc(1) | 4118.4495 |
| HexNAc(8)Hex(9)Fuc(1)NeuAc(1)NeuGc(2) | 4134.4444 |
| HexNAc(11)Hex(11)NeuAc(1) | 4307.5496 |
| HexNAc(11)Hex(11)NeuGc(1) | 4323.5445 |
| HexNAc(8)Hex(9)Fuc(1)NeuAc(4) | 4393.5500 |
| HexNAc(9)Hex(10)Fuc(1)NeuAc(4) | 4758.6822 |

**Supplementary Table S4**. Glycan search space (200 N-glycan compositions) used for the Byonic searches of human sera for the on-line proof-of-concept experiments. Masses are provided as neutral monoisotopic values (M).

| **Glycan composition** | **Mass (Da)** |
|---|---|
| HexNAc(1) | 203.0794 |
| HexNAc(2) | 406.1587 |
| HexNAc(2)Hex(1) | 568.2116 |
| HexNAc(2)Hex(2) | 730.2644 |
| HexNAc(1)Fuc(1) | 349.1373 |
| HexNAc(2)Fuc(1) | 552.2167 |
| HexNAc(2)Hex(1)Fuc(1) | 714.2695 |
| HexNAc(2)Hex(2)Fuc(1) | 876.3223 |
| HexNAc(2)Hex(3)Fuc(1) | 1038.3750 |
| HexNAc(2)Hex(3) | 892.3172 |
| HexNAc(2)Hex(4) | 1054.3700 |
| HexNAc(3)Hex(3) | 1095.3966 |
| HexNAc(2)Hex(3)Fuc(2) | 1184.4330 |
| HexNAc(2)Hex(4)Fuc(1) | 1200.4280 |
| HexNAc(2)Hex(5) | 1216.4229 |
| HexNAc(3)Hex(3)Fuc(1) | 1241.4545 |
| HexNAc(3)Hex(4) | 1257.4494 |
| HexNAc(4)Hex(3) | 1298.4760 |
| HexNAc(2)Hex(5)Fuc(1) | 1362.4808 |
| HexNAc(2)Hex(6) | 1378.4757 |
| HexNAc(3)Hex(3)Fuc(2) | 1387.5124 |
| HexNAc(3)Hex(4)Fuc(1) | 1403.5073 |
| HexNAc(3)Hex(5) | 1419.5022 |
| HexNAc(4)Hex(3)Fuc(1) | 1444.5339 |
| HexNAc(4)Hex(4) | 1460.5288 |
| HexNAc(5)Hex(3) | 1501.5553 |
| HexNAc(2)Hex(7) | 1540.5285 |
| HexNAc(3)Hex(4)NeuAc(1) | 1548.5448 |
| HexNAc(3)Hex(4)Fuc(2) | 1549.5652 |
| HexNAc(3)Hex(5)Fuc(1) | 1565.5601 |
| HexNAc(3)Hex(6) | 1581.5551 |
| HexNAc(4)Hex(3)NeuAc(1) | 1589.5714 |
| HexNAc(4)Hex(3)Fuc(2) | 1590.5918 |
| HexNAc(4)Hex(4)Fuc(1) | 1606.5867 |
| HexNAc(4)Hex(5) | 1622.5816 |
| HexNAc(5)Hex(3)Fuc(1) | 1647.6132 |
| HexNAc(5)Hex(4) | 1663.6082 |
| HexNAc(3)Hex(4)Fuc(1)NeuAc(1) | 1694.6027 |
| HexNAc(2)Hex(8) | 1702.5813 |
| HexNAc(6)Hex(3) | 1704.6347 |
| HexNAc(3)Hex(5)NeuAc(1) | 1710.5977 |
| HexNAc(3)Hex(6)Fuc(1) | 1727.6130 |
| HexNAc(4)Hex(3)Fuc(3) | 1736.6497 |
| HexNAc(4)Hex(4)NeuAc(1) | 1751.6242 |
| HexNAc(4)Hex(4)Fuc(2) | 1752.6446 |
| HexNAc(4)Hex(5)Fuc(1) | 1768.6395 |
| HexNAc(4)Hex(6) | 1784.6344 |
| HexNAc(5)Hex(4)Fuc(1) | 1809.6661 |
| HexNAc(5)Hex(5) | 1825.6610 |
| HexNAc(6)Hex(3)Fuc(1) | 1850.6926 |
| HexNAc(2)Hex(9) | 1864.6342 |
| HexNAc(6)Hex(4) | 1866.6875 |
| HexNAc(3)Hex(6)NeuAc(1) | 1872.6505 |
| HexNAc(4)Hex(4)Fuc(1)NeuAc(1) | 1897.6821 |
| HexNAc(7)Hex(3) | 1907.7141 |
| HexNAc(4)Hex(5)NeuAc(1) | 1913.6770 |
| HexNAc(4)Hex(5)Fuc(2) | 1914.6974 |
| HexNAc(4)Hex(6)Fuc(1) | 1930.6923 |
| HexNAc(5)Hex(3)Fuc(1)NeuAc(1) | 1938.7087 |
| HexNAc(4)Hex(7) | 1946.6873 |
| HexNAc(5)Hex(4)NeuAc(1) | 1954.7036 |
| HexNAc(5)Hex(4)Fuc(2) | 1955.7240 |
| HexNAc(5)Hex(5)Fuc(1) | 1971.7189 |
| HexNAc(5)Hex(6) | 1987.7138 |
| HexNAc(6)Hex(3)Fuc(2) | 1996.7505 |
| HexNAc(6)Hex(4)Fuc(1) | 2012.7454 |
| HexNAc(3)Hex(6)Fuc(1)NeuAc(1) | 2018.7084 |
| HexNAc(2)Hex(10) | 2026.6870 |
| HexNAc(6)Hex(5) | 2028.7404 |
| HexNAc(4)Hex(4)Fuc(2)NeuAc(1) | 2043.7400 |
| HexNAc(7)Hex(3)Fuc(1) | 2053.7720 |
| HexNAc(4)Hex(5)Fuc(1)NeuAc(1) | 2059.7349 |
| HexNAc(4)Hex(5)Fuc(3) | 2060.7553 |

| | |
|---|---|
| HexNAc(7)Hex(4) | 2069.7669 |
| HexNAc(4)Hex(6)NeuAc(1) | 2075.7299 |
| HexNAc(4)Hex(6)Fuc(2) | 2076.7503 |
| HexNAc(4)Hex(7)Fuc(1) | 2092.7452 |
| HexNAc(5)Hex(4)Fuc(1)NeuAc(1) | 2100.7615 |
| HexNAc(8)Hex(3) | 2110.7935 |
| HexNAc(5)Hex(5)NeuAc(1) | 2116.7564 |
| HexNAc(5)Hex(5)Fuc(2) | 2117.7768 |
| HexNAc(5)Hex(6)Fuc(1) | 2133.7717 |
| HexNAc(6)Hex(3)Fuc(1)NeuAc(1) | 2141.7880 |
| HexNAc(6)Hex(3)Fuc(3) | 2142.8084 |
| HexNAc(5)Hex(7) | 2149.7666 |
| HexNAc(6)Hex(4)NeuAc(1) | 2157.7830 |
| HexNAc(6)Hex(4)Fuc(2) | 2158.8034 |
| HexNAc(6)Hex(5)Fuc(1) | 2174.7983 |
| HexNAc(2)Hex(11) | 2188.7398 |
| HexNAc(6)Hex(6) | 2190.7932 |
| HexNAc(4)Hex(5)NeuAc(2) | 2204.7724 |
| HexNAc(4)Hex(5)Fuc(2)NeuAc(1) | 2205.7928 |
| HexNAc(4)Hex(5)Fuc(4) | 2206.8132 |
| HexNAc(7)Hex(4)Fuc(1) | 2215.8248 |
| HexNAc(4)Hex(6)Fuc(1)NeuAc(1) | 2221.7878 |
| HexNAc(4)Hex(6)Fuc(3) | 2222.8082 |
| HexNAc(4)Hex(7)NeuAc(1) | 2237.7827 |
| HexNAc(4)Hex(7)Fuc(2) | 2238.8031 |
| HexNAc(5)Hex(4)NeuAc(2) | 2245.7990 |
| HexNAc(5)Hex(4)Fuc(2)NeuAc(1) | 2246.8194 |
| HexNAc(8)Hex(3)Fuc(1) | 2256.8514 |
| HexNAc(5)Hex(5)Fuc(1)NeuAc(1) | 2262.8143 |
| HexNAc(5)Hex(5)Fuc(3) | 2263.8347 |
| HexNAc(8)Hex(4) | 2272.8463 |
| HexNAc(5)Hex(6)NeuAc(1) | 2278.8092 |
| HexNAc(5)Hex(6)Fuc(2) | 2279.8296 |
| HexNAc(6)Hex(3)Fuc(2)NeuAc(1) | 2287.8459 |
| HexNAc(5)Hex(7)Fuc(1) | 2295.8245 |
| HexNAc(5)Hex(8) | 2311.8195 |
| HexNAc(9)Hex(3) | 2313.8728 |
| HexNAc(6)Hex(5)Fuc(2) | 2320.8562 |
| HexNAc(6)Hex(6)Fuc(1) | 2336.8511 |
| HexNAc(2)Hex(12) | 2350.7926 |
| HexNAc(4)Hex(5)Fuc(1)NeuAc(2) | 2350.8304 |
| HexNAc(6)Hex(7) | 2352.8460 |

| | |
|---|---|
| HexNAc(7)Hex(4)Fuc(2) | 2361.8827 |
| HexNAc(5)Hex(4)Fuc(1)NeuAc(2) | 2391.8569 |
| HexNAc(7)Hex(6) | 2393.8726 |
| HexNAc(5)Hex(5)NeuAc(2) | 2407.8518 |
| HexNAc(5)Hex(6)Fuc(1)NeuAc(1) | 2424.8671 |
| HexNAc(5)Hex(6)Fuc(3) | 2425.8875 |
| HexNAc(6)Hex(3)Fuc(1)NeuAc(2) | 2432.8835 |
| HexNAc(8)Hex(5) | 2434.8991 |
| HexNAc(5)Hex(8)Fuc(1) | 2457.8774 |
| HexNAc(9)Hex(3)Fuc(1) | 2459.9307 |
| HexNAc(9)Hex(4) | 2475.9257 |
| HexNAc(6)Hex(6)Fuc(2) | 2482.9090 |
| HexNAc(6)Hex(7)Fuc(1) | 2498.9039 |
| HexNAc(7)Hex(6)Fuc(1) | 2539.9305 |
| HexNAc(5)Hex(5)Fuc(1)NeuAc(2) | 2553.9097 |
| HexNAc(7)Hex(7) | 2555.9254 |
| HexNAc(5)Hex(6)NeuAc(2) | 2569.9046 |
| HexNAc(5)Hex(6)Fuc(2)NeuAc(1) | 2570.9250 |
| HexNAc(8)Hex(5)Fuc(1) | 2580.9570 |
| HexNAc(5)Hex(7)Fuc(1)NeuAc(1) | 2586.9200 |
| HexNAc(8)Hex(6) | 2596.9519 |
| HexNAc(5)Hex(9)Fuc(1) | 2619.9302 |
| HexNAc(9)Hex(4)Fuc(1) | 2621.9836 |
| HexNAc(6)Hex(6)Fuc(3) | 2628.9669 |
| HexNAc(4)Hex(5)Fuc(3)NeuAc(2) | 2642.9462 |
| HexNAc(4)Hex(5)Fuc(3)NeuAc(1) | 2351.8508 |
| HexNAc(6)Hex(7)NeuAc(1) | 2643.9414 |
| HexNAc(6)Hex(9) | 2676.9517 |
| HexNAc(7)Hex(7)Fuc(1) | 2701.9833 |
| HexNAc(5)Hex(6)Fuc(1)NeuAc(2) | 2715.9626 |
| HexNAc(5)Hex(6)Fuc(3)NeuAc(1) | 2716.9830 |
| HexNAc(7)Hex(8) | 2717.9782 |
| HexNAc(8)Hex(7) | 2759.0048 |
| HexNAc(6)Hex(6)NeuAc(1) | 2481.8886 |
| HexNAc(6)Hex(6)NeuAc(2) | 2772.9840 |
| HexNAc(6)Hex(7)Fuc(1)NeuAc(1) | 2789.9993 |
| HexNAc(9)Hex(6) | 2800.0313 |
| HexNAc(6)Hex(8)NeuAc(1) | 2805.9942 |
| HexNAc(5)Hex(6)NeuAc(3) | 2861.0001 |
| HexNAc(7)Hex(8)Fuc(1) | 2864.0361 |
| HexNAc(5)Hex(7)Fuc(1)NeuAc(2) | 2878.0154 |
| HexNAc(5)Hex(8)Fuc(4) | 2896.0511 |

| | |
|---|---|
| HexNAc(6)Hex(6)Fuc(1)NeuAc(2) | 2919.0419 |
| HexNAc(6)Hex(6)Fuc(1)NeuAc(1) | 2627.9465 |
| HexNAc(8)Hex(8) | 2921.0576 |
| HexNAc(6)Hex(7)NeuAc(2) | 2935.0368 |
| HexNAc(6)Hex(7)Fuc(4) | 2937.0776 |
| HexNAc(9)Hex(6)Fuc(1) | 2946.0892 |
| HexNAc(6)Hex(8)Fuc(1)NeuAc(1) | 2952.0522 |
| HexNAc(5)Hex(6)Fuc(1)NeuAc(3) | 3007.0580 |
| HexNAc(7)Hex(8)NeuAc(1) | 3009.0736 |
| HexNAc(6)Hex(5)Fuc(1)NeuAc(2) | 2756.9891 |
| HexNAc(6)Hex(5)Fuc(1)NeuAc(1) | 2465.8937 |
| HexNAc(6)Hex(5)Fuc(1)NeuAc(3) | 3048.0845 |
| HexNAc(8)Hex(8)Fuc(1) | 3067.1155 |
| HexNAc(6)Hex(7)Fuc(1)NeuAc(2) | 3081.0947 |
| HexNAc(8)Hex(9) | 3083.1104 |
| HexNAc(6)Hex(7)Fuc(5) | 3083.1356 |
| HexNAc(6)Hex(11)Fuc(1) | 3147.1152 |
| HexNAc(10)Hex(7) | 3165.1635 |
| HexNAc(6)Hex(6)Fuc(1)NeuAc(3) | 3210.1373 |
| HexNAc(6)Hex(7)NeuAc(3) | 3226.1323 |
| HexNAc(6)Hex(7)Fuc(4)NeuAc(1) | 3228.1731 |
| HexNAc(8)Hex(9)Fuc(1) | 3229.1683 |
| HexNAc(6)Hex(10)Fuc(1)NeuAc(1) | 3276.1578 |

| | |
|---|---|
| HexNAc(7)Hex(7)Fuc(1)NeuAc(2) | 3284.1741 |
| HexNAc(6)Hex(7)Fuc(1)NeuAc(3) | 3372.1902 |
| HexNAc(6)Hex(9)Fuc(1)NeuAc(2) | 3405.2004 |
| HexNAc(9)Hex(9)Fuc(1) | 3432.2477 |
| HexNAc(7)Hex(8)Fuc(1)NeuAc(1) | 3155.1315 |
| HexNAc(7)Hex(8)Fuc(1)NeuAc(2) | 3446.2269 |
| HexNAc(9)Hex(10) | 3448.2426 |
| HexNAc(6)Hex(7)NeuAc(4) | 3517.2277 |
| HexNAc(7)Hex(7)Fuc(1)NeuAc(3) | 3575.2695 |
| HexNAc(9)Hex(10)Fuc(1) | 3594.3005 |
| HexNAc(6)Hex(7)Fuc(1)NeuAc(4) | 3663.2856 |
| HexNAc(7)Hex(6)Fuc(1)NeuAc(4) | 3704.3121 |
| HexNAc(7)Hex(8)Fuc(1)NeuAc(3) | 3737.3224 |
| HexNAc(10)Hex(10)Fuc(1) | 3797.3799 |
| HexNAc(7)Hex(7)Fuc(1)NeuAc(4) | 3866.3650 |
| HexNAc(7)Hex(8)Fuc(1)NeuAc(4) | 4028.4178 |
| HexNAc(8)Hex(9)Fuc(1)NeuAc(3) | 4102.4546 |
| HexNAc(11)Hex(11)NeuAc(1) | 4307.5496 |
| HexNAc(8)Hex(9)Fuc(1)NeuAc(4) | 4393.5500 |
| HexNAc(9)Hex(10)Fuc(1)NeuAc(4) | 4758.6822 |

**References used in Supplemental Information**


1 Dey, K. K. *et al.* Deep undepleted human serum proteome profiling toward biomarker discovery for Alzheimer's disease. *Clin Proteomics* **16**, 16 (2019). https://doi.org/10.1186/s12014-019-9237-1

2 Kawahara, R. *et al.* Multi-omics definition of the sex-specific glycoproteome of murine tissues. *bioRxiv*, 2026.2003.2010.710926 (2026). https://doi.org/10.64898/2026.03.10.710926
